# PPARγ is a tumor suppressor in basal bladder tumors offering new potential therapeutic opportunities

**DOI:** 10.1101/868190

**Authors:** Laure Coutos-Thévenot, Syrine Beji, Hélène Neyret-Kahn, Quentin Pippo, Jacqueline Fontugne, Judith Osz, Clémentine Krucker, Clarice Dos Santos Groeneveld, Florent Dufour, Aurélie Kamoun, Marie Ley, Elodie Chapeaublanc, Aurélien de Reynies, Thierry Lebret, Yves Allory, Sarah Cianférani, François Radvanyi, Natacha Rochel, Isabelle Bernard-Pierrot

## Abstract

PPARγ activation is a critical event in luminal muscle-invasive bladder cancer (MIBC) tumorigenesis, favoring both tumor cell growth and microenvironment modulation toward tumor immune escape. Conversely, the down-regulation of PPARγ activity in basal MIBC suggests tumor suppressive effects in this subgroup. Here, we report genetic, epigenetic and functional evidence to support the tumor suppressor role for *PPARγ* in basal bladder tumors. We identified hemizygous deletions, DNA hyper-methylation and loss-of-function mutations of PPARγ in basal MIBC, associated with *PPARγ* under-expression and its decreased activity. Re-expression of PPARγ in basal tumor cells resulted in the activation of PPARγ -dependent transcription program that modulated fatty acid metabolism and cell differentiation and decreased cell growth, which could partly rely on EGFR down-regulation. Structure-function studies of two PPARγ mutant revealed a destabilization of a region important for coactivator recruitment and should help develop potent molecules to activate PPARγ as a therapeutic strategy for basal MIBC. The identification of this subtype-dependent dual role of PPARγ in MIBC strengthens the critical role of PPARγ in bladder tumorigenesis and reinforces the interest in stratified medicine based on tumor molecular subtyping.

**One sentence summary:** Genetic, epigenetic and functional evidence of a tumor suppressor role for *PPARγ* in basal bladder tumors offer new therapeutic opportunities for this subgroup.

## Introduction

The nuclear receptor PPARγ (peroxisome proliferator-activated receptor gamma) functions as a permissive heterodimer with RXRα (retinoid X receptor) and recognizes specific sequence motifs, defined as PPRE (peroxisome proliferative response elements), in the regulatory regions of its target genes. In the absence of ligand, PPARγ is complexed with corepressor proteins, such as NCoR1 (Nuclear receptor corepressor 1) or SMRT (silencing mediator of retinoic acid receptor), which induces HDAC (histone deacetylase) recruitment, leading to PPARγ-mediated transcriptional repression. Conversely, upon ligand binding, a conformational change allows the release of corepressors and the recruitment of coactivators, such as MED1 (Mediator complex subunit 1) or PGC1α (PPARGC1A, PPARG coactivator 1 alpha), enabling PPARγ transcriptional activity. A variety of natural ligands, such as polyunsaturated fatty acid or prostaglandin J2 derivatives, and synthetic ligands, such as thiazolinediones, can activate PPARγ. PPARγ is involved in the regulation of glucose homeostasis and adipogenesis^1,2^ but also in the differentiation of several tissue types, including the urothelium^3,4^. Its role in cancer is less clear and seems to be dual, tumor suppressor (in colon, lung cancers and neuroblastoma) or pro-tumorigenic (in pancreatic or bladder cancers), depending on the cell type^5–9^.

In bladder cancer, the 4^th^ most frequent cancer in men in industrialized countries, the luminal subtype of muscle-invasive bladder carcinomas (MIBCs), which accounts for 60% of MIBCs^10,11^, has been shown to display a PPARγ activation signature^8,12^. This activation is associated with genetic alterations which are associated with the luminal subtype, namely PPARγ DNA gains and amplifications (30%) or recurrent activating mutations of RXRα (5%) or PPARγ (4%)^10,13–16^. PPARγ activation renders bladder tumor cell growth PPARγ-dependent^14,16^ and promotes immune evasion in MIBCs^17^ providing evidence for pro-tumorigenic roles of the PPARγ/RXRα pathway in luminal bladder tumors.

Interestingly, PPARγ activation signature or regulon activity is dramatically decreased in basal bladder tumors, a subtype accounting for 35% of MIBCs and presenting a poor prognosis, suggesting that PPARγ could display an opposite role in this subtype^10–12,18^. In this work, we hypothesized that the loss of PPARγ activity could be essential for the tumorigenesis of the basal subtype of MIBCs. We searched for genetic and epigenetic alterations that could drive PPARγ inactivation in basal bladder tumors and could support a tumor suppressor role for PPARγ. In basal tumors, we identified an enrichment in hemizygous deletions, DNA hyper-methylation and repressive histone mark (H3K9me3) of PPARγ, associated with PPARγ loss of expression. Among the non-recurrent mutations of PPARγ that we previously identified by sequencing PPARγ in 359 tumors and studying publicly available data for 455 MIBCs^16^, four mutations were associated with basal tumors. Functional analysis revealed that these four mutations reduce the transcriptional activity of PPARγ. Further biochemical and structure-function analysis of two of these mutations, affecting the ligand-binding domain of PPARγ, showed that they alter PPARγ activity through the destabilization of helix H12, thereby impairing the release of corepressors and the recruitment of coactivators. Furthermore, induced PPARγ expression in basal bladder cancer cell lines activated PPARγ-dependent transcription and decreased cell viability, whereas it displayed no effect on cell viability of luminal cells. Finally, we show that the tumor suppressive activity of PPARγ in basal tumors could at least partially rely on EGFR down-regulation. Our study provides genetic, epigenetic and functional evidence of a tumor suppressive role for PPARγ in basal bladder cancer and therefore supports the use of PPARγ synthetic agonists as a therapeutic strategy for this subgroup. It reinforces the central role of PPARγ in bladder tumorigenesis that appears to be subgroup dependent: pro-tumorigenic in differentiated luminal tumors and tumor suppressor in basal tumors.

## Results

### Hemizygous deletions, DNA hypermethylation and loss of expression of PPARγ associated with basal MIBC

We studied *PPARγ* expression in 197 bladder tumors −101 of which were Non Muscle-Invasive Bladder Cancers (NMIBCs)-in our CIT series of tumors (Carte d’Identité des Tumeurs) using U133 plus 2.0 Affymetrix transcriptomic data^8,18^ and in 405 MIBC samples using publicly available RNAseq data from The Cancer Genome Atlas^10,13^ genomic database (http://cancergenome.nih.gov) (Fig. 1a). Tumors were grouped in six molecular classes according to a molecular consensus classification derived from six independent classification systems previously described for MIBC^11^ (Supplementary Table 1). In good agreement with the loss of transcriptional activity of PPARγ observed in basal tumors^10–12^, we observed a significantly lower expression of PPARγ in Basal/Squamous tumors but overall in non-luminal tumors including in Neuroendocrine-like and Stroma-rich tumors compared to luminal tumors, encompassing Luminal Papillary, Luminal Non-Specified and Luminal Unstable subtypes (Fig.1a). Using available copy number data for 385 MIBCs from the TCGA, we observed that hemizygous deletions of PPARγ were significantly associated with a low expression (Fig.1b left panel) and activity (Fig.1b right panel) of PPARγ. Conversely, gains and amplifications were associated with overexpression and hyper-activity of PPARγ, as previously described^8,9,14,16^. We further noticed that these hemizygous deletions were significantly enriched in basal/squamous tumors and could therefore account for part of the loss of expression of PPARγ in this subgroup (Fig. 1c). The genomic deletions of PPARγ in basal tumors were a first genomic evidence suggesting a tumor suppressor role for PPARγ in these tumors. Since aberrant patterns of DNA methylation can also affect tumor suppressor gene expression during tumorigenesis, we analyzed PPARγ DNA methylation in 368 MIBCs using DNA methylation array data available from TCGA. We identified a significant hyper-methylation of PPARγ CpGs in basal tumors compared to luminal ones, mostly in shore regions of CpG islands within the PPARγ promoter (Fig.1d, upper panel and supplementary Fig.1a). The hyper-methylation, which correlated with the downregulation of PPARγ expression in basal tumors (Fig.1d, middle and lower panels), could also account for part of its loss of expression in basal tumors. These results were further validated in our CIT series of tumors (Supplementary Fig.1b). We also observed an enrichment of histone repressive mark (H3K9me3) in PPARγ regulatory regions in two basal (L1207 and 5367) as compared to two luminal bladder cancer cell lines (RT112 and SD48) (Supplementary Fig.1c). These data provided epigenetic evidences supporting the tumor suppressor role of PPARγ in basal tumors.

**Figure 1:**
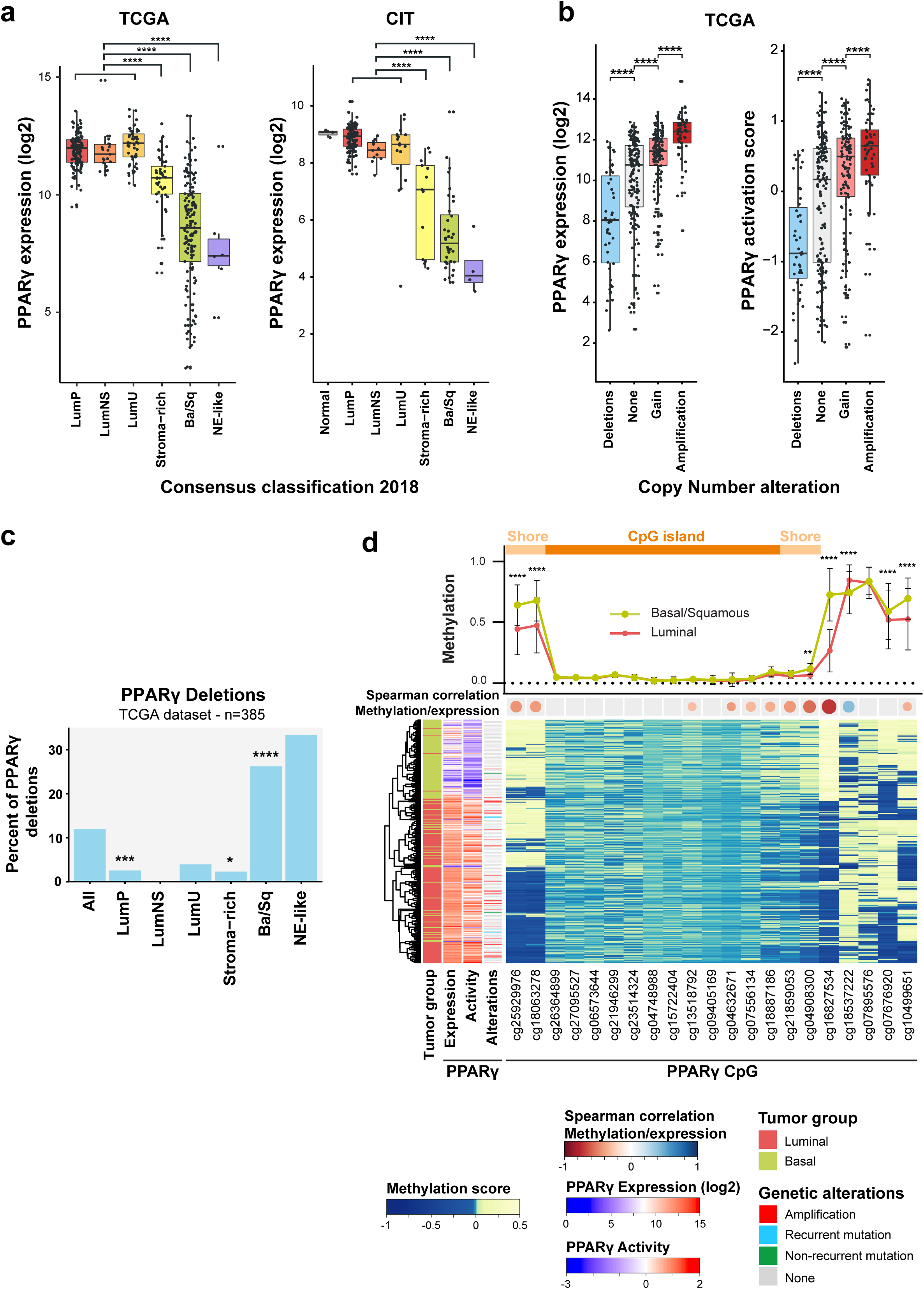
Hemizygous deletions and DNA hypermethylation are associated with loss of expression of PPARγ in basal bladder tumors. **a)** Expression level of PPARγ in 336 bladder tumors from our CIT series of tumors (Affymetrix U133 plus2.0 signal) and in 405 MIBCs from TCGA (RNA-seq). Tumors were classified according to a consensus molecular classification that defines six subgroups of MIBC (Kamoun *et al.*, 2018). PPARγ expression levels in stroma-rich, basal/squamous (Ba/Sq) and in Neuroendocrine-like (NE-like) tumors were compared to those in luminal tumors, comprising luminal papillary (LumP), Luminal non specified (LumNS) and Luminal unstable (LumU) tumors using Dunnett’s multiple comparison test, **** *p*<0.0001. **b)** Expression level and activity of PPARγ were compared in relation to PPARγ copy number alterations in 385 MIBCs from TCGA using Dunnett’s multiple comparison test, **** *p*<0.0001.**c)** Frequency of genomic heterozygous deletion of PPARγ in the different consensus subgroups of MIBC. Enrichment or depletion in genomic PPARγ deletion in the different subgroups were evaluated using Fisher exact t-test, * 0.01<*p*<0.05; *** 0.0001<p< 0.001; **** *p*<0.0001.**d)** Methylation of each CpG in TCGA dataset for basal (green) and luminal (red) tumors (upper panel), correlation of CpG methylation and PPARγ expression (middle panel) and heatmap representing centered methylation score for each tumor (lower panel). Enrichment in CpG methylation in the different subgroups were evaluated using 2way ANOVA test, * 0.01<*p*<0.05; *** 0.0001<p< 0.001; **** *p*<0.0001

### Loss-of-function mutations of PPARG in basal tumors

Loss-of-function mutations are also a hallmark of tumor suppressor genes. We therefore searched for such alterations of PPARγ in basal bladder tumors. We previously sequenced PPARγ in 359 bladder tumors and studied publicly available data for 455 MIBCs, which allowed us to identify recurrent activating mutations of PPARγ in 4% of bladder tumors^16^. These mutations were enriched in tumors presenting a high PPARγ activation score, which were mostly luminal tumors, supporting the protumorigenic role of PPARγ in this subgroup. We had also identified eleven non-recurrent PPARγ mutations that we did not further study^16^ (Fig. 2a). Here, we numbered all mutations relative to the PPARγ*2* isoform (NM_015869), which is 28 amino acids longer than the PPARγ*1* isoform (NM_138712) at the N-terminal end (Fig. 2a). Four of these non-recurrent mutations, S74C, F310S, E455Q and H494Y, were associated with basal tumors which presented a low PPARγ activation score (Fig. 2b). We therefore hypothesized that these four mutations could be loss-of-function mutations and investigated their functional impact on the transcriptional activity of PPARγ (Fig. 2c and 2d). We used, in HEK293FT cells, a luciferase reporter gene containing three copies of the DR1 sequence of the PPRE arranged in tandem and linked to the thymidine kinase promoter (PPRE-3X-TK)^19^. The four PPARγ mutant proteins had significantly lower levels of transcriptional activity than the wild type even in the presence of rosiglitazone, a synthetic PPARγ agonist (activity reduced by 25% to 90%) (Fig. 2c). We further focused on the two most inactive mutants that affect the ligand binding domain (LBD) of PPARγ, F310S and H494Y. We showed that their overexpression in the basal bladder cell line 5637 induced a significantly lower expression of several known PPARγ target genes (*FABP4* and *ACSL5*) compared to the wild-type protein as shown by RT-qPCR (Fig. 2d). The effects of the mutants were comparable to that of the empty plasmid, suggesting an absence of transcriptional activity of the proteins in this system. The results of these two different approaches to measure PPARγ transcriptional activity clearly showed that PPARγ mutations F310S and H494Y are loss-of-function mutations affecting basal tumors.

**Figure 2:**
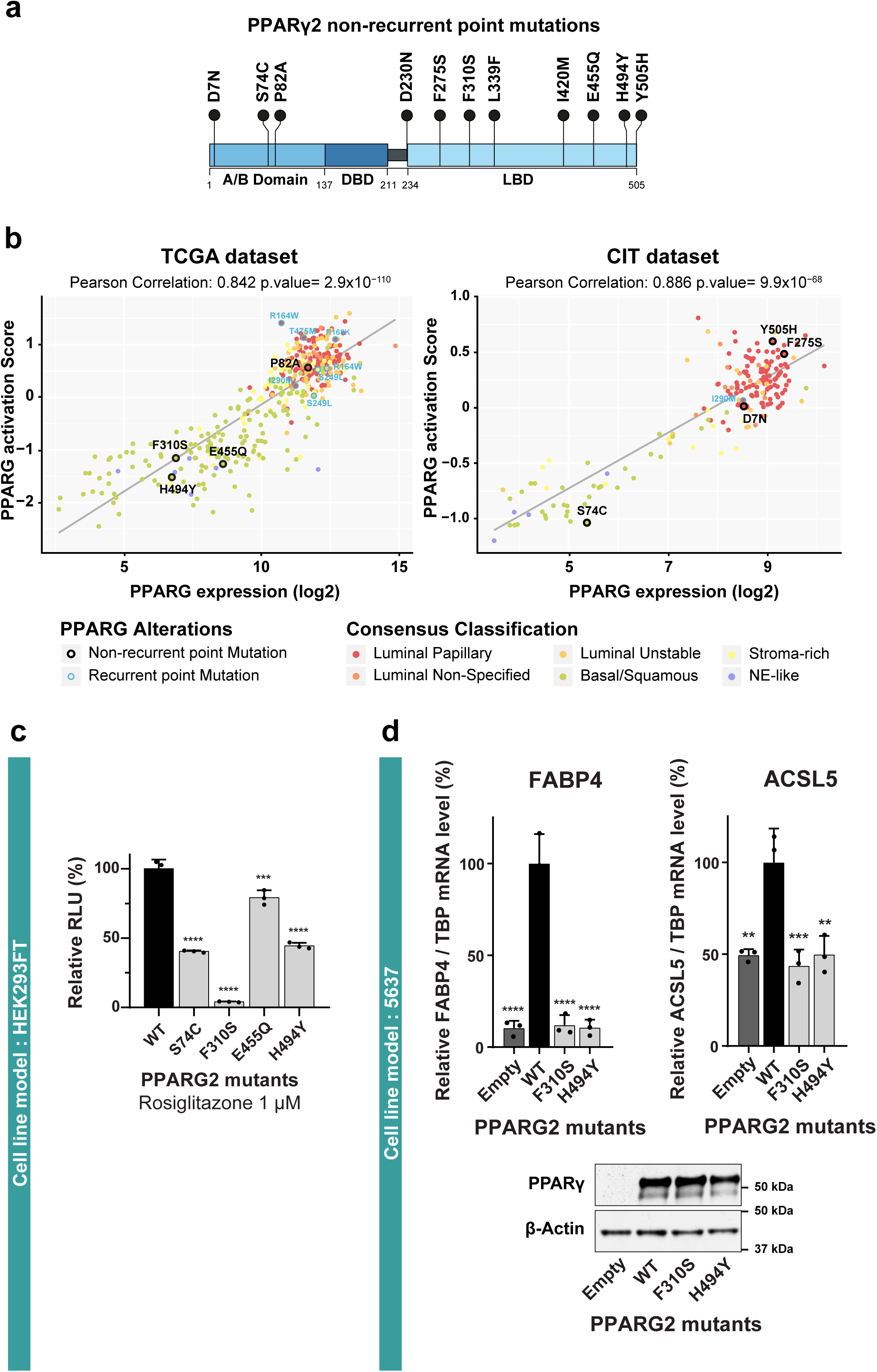
Transcriptional activity of non-recurrent PPARγ mutants identified in basal tumors. **a)** Lolliplot representation of non-recurrent mutations of PPARγ that we previously identified in 859 bladder tumors (Rochel et al., 2019). Sequences are numbered according to the PPARγ2 isoform. A/B: N-terminal domain; DBD: DNA-binding domain; LBD: ligand-binding domain. **b)** PPARγ expression levels and activation scores in 403 and 196 tumors from TCGA and CIT series respectively, for which both PPARγ mutation data and transcriptomic data were available. **c)** A reporter plasmid containing the firefly luciferase gene under the control of a PPRE-X3-TK promoter was co-expressed in HEK293FT cells with a pcDNA3 vector encoding wild-type (WT) or mutant PPARγ2 associated with basal tumors (S74C, F310S, E455Q, H494Y). Cells were stimulated with 1µM rosiglitazone. *Renilla* luciferase, expressed under the control of the CMV promoter, was used to normalize the signal. The data shown are the means ± SD of one representative experiment conducted in triplicate. The results for each mutant were compared with those for the wild-type using Dunnett’s multiple comparison test, *** 0.0001<*p*<0.001; **** *p*<0.0001 **d)** 5637 basal cells were transiently transfected with a pcDNA3 vector encoding wild-type (WT) or mutant of the LBD domain (F310S, H494Y) PPARγ. The expression of all PPARγ forms was checked by western blotting, β-actin was used as loading control (lower panel). The expression of two PPARγ target genes was normalized against TBP expression and is shown as percentage relative to the expression induced by wild-type PPARγ (upper panel). The data are presented as the mean ± SD of three independent experiments. The results for each mutant were compared with those for the wild type using Dunnett’s multiple comparison test: ** 0.001<*p*<0.01;*** 0.0001<p< 0.001; **** *p*<0.0001.

### Loss-of-function mutations impair the release of corepressors and recruitment of coactivators by PPARγ in the presence of ligand

We then performed biochemical and biophysical analyses to understand how these two mutations impair PPARγ activity. F310S is located in the N-ter of helix 3 facing the loop between helices 11 and 12 whereas the H494Y mutation involves a residue at the N-terminal boundary of helix 12 (Fig. 3a). Native electrospray mass spectrometry indicated that the purified recombinant LBD wild-type and mutants (Supplementary Figs. 2-3) were not bound to any ligand (Supplementary Fig. 4). We used nano differential scanning fluorimetry to compare the thermal stability of the purified PPARγ WT and mutants, alone and upon binding to the agonist ligand, GW1929 (Supplementary Fig. 2c). The two mutants in their apo form exhibited a lower melting temperature (Tm) than the WT with a ΔTm of 1°C and 2°C for F310S and H494Y, respectively. Of note, GW1929 induced a strong stabilization of the WT, as well as of the 2 mutants, suggesting that the 2 mutations do not significantly affect the binding of GW1929. Both mutants efficiently formed heterodimers with RXRα similarly to PPARγ wild-type (Supplementary Fig. 2b). We therefore focused on the ability of mutations to modulate PPARγ binding to corepressors and coactivators, which involves recognition of motifs on coregulators by the LBD, and ultimately regulates the transcriptional output of target genes. To study the binding of PPARγ to corepressor and coactivator proteins, we performed mammalian two-hybrid assay in HEK293FT cells, using VP16-fused PPARγ (wild-type, F310S and H494Y), GAL4-DNA-binding-domain-fused co-repressor (NCoR1 or SMRT) or co-activator MED1 and pG5-LUC reporter. We showed that in the context of full protein and in presence of exogenous ligand (rosiglitazone), the two mutations significantly favored the binding of the two co-repressor peptides and inhibited the recruitment of MED1 coactivator domain compared to PPARγ wild-type (Fig. 3b). The lower recruitment of the coactivator and increased recruitment of the corepressors by the two mutants was confirmed by monitoring coregulator peptide recruitment by the different LBDs (Fig.3c). We measured the interaction between wild-type or mutant forms of PPARγ and a fluorescently labeled coactivator peptide of PGC1α (PPARGC1A) or a fluorescent labeled NCoR1 corepressor peptide, by MicroScale Thermophoresis. In the absence of ligand, the WT recruited the coactivator peptide with higher affinity than the two mutants (Fig. 3c). The addition of a full agonist, rosiglitazone (Supplementary Fig. 5), enhanced the interaction between PPARγ and PGC1α coactivator peptide, with the WT exhibiting again the highest affinity. On the other hand, the 2 mutants exhibited increased affinity for the corepressor peptide of NCoR1 compared to WT (Fig. 3c). The increased interaction with corepressor and decreased interaction with coactivator of the mutants was also observed by native mass spectrometry (Supplementary Figs. 6-8). Together, these data suggest that the two considered mutations, F310S and H494Y, impair the adoption of an agonist conformation by PPARγ in the presence of ligand, thereby enhancing corepressor interactions and inhibiting coactivator interaction.

**Figure 3:**
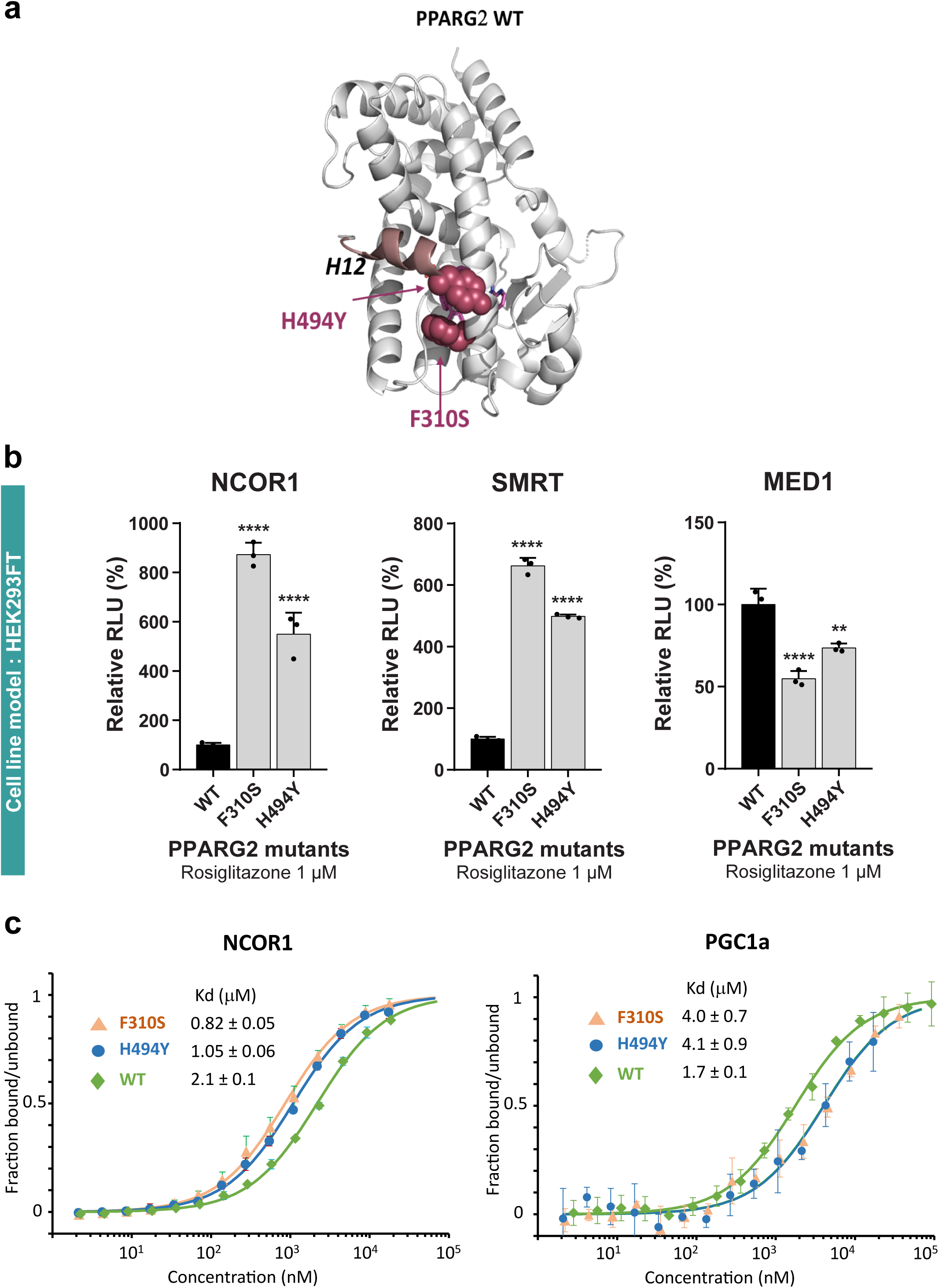
Effect of PPARγ F310S and H494Y mutations on coregulator interactions. **a)** Position of the residues affected by non-recurrent PPARγ mutations associated with basal tumors on the 3D structure of PPARγ LBD. **b)** Mammalian two-hybrid analysis in HEK293T cells. pG5-Firefly luciferase reporter plasmid was co-expressed with VP16-PPARG (wild-type or mutant full-length proteins) and with GAL4-DNA-binding-domain-fused NCoR1 or SMRT corepressor or MED1 coactivator. *Renilla* luciferase, expressed under the control of the CMV promoter, was used to normalize the signal. The data shown are the means ± SD of one representative experiment conducted in quadruplicate. The results for each mutant were compared with those for the wild-type using Dunnett’s multiple comparison test, ** 0.001< *p*<0.01, **** p<0,0001. **c)** Effect of PPARγ mutations on NCoR1 peptide (left) and PGC1α peptide (right) interactions as determined by microscale thermophoresis. Unlabeled PPARγ LBD protein was titrated into a fixed concentration of fluorescently labeled peptide in the absence of ligand. Isotherms were averaged over three consecutive measurements ^and fitted according to the law of mass action to yield the apparent K^d^. Each plot is^ representative of at least two independent experiments performed with different batches of protein preparation.

**Figure 4:**
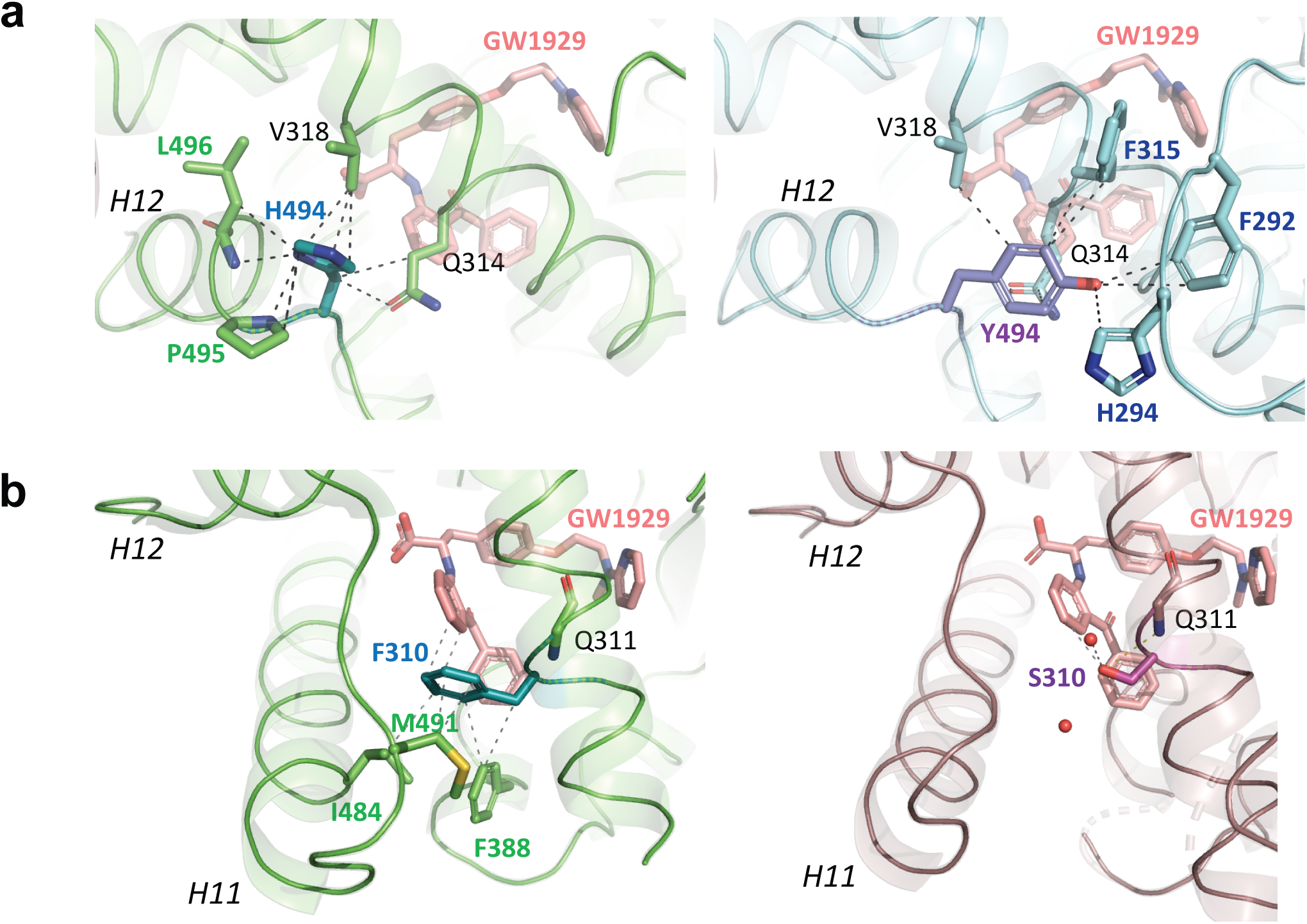
Impact of PPARγ F310S and H494Y mutations on the crystal structure of the protein. **a)** Close-up of the regions around the H494Y mutation, showing its interactions in the WT complex (left) and in the mutant complex (right). PPARγ WT and H494Y are in green and cyan, respectively, with the coactivator peptide in plum. **b)** Close-up of the regions around the F310S mutation, showing its interactions in the WT complex (left) and in the mutant complex (right).

**Figure 5:**
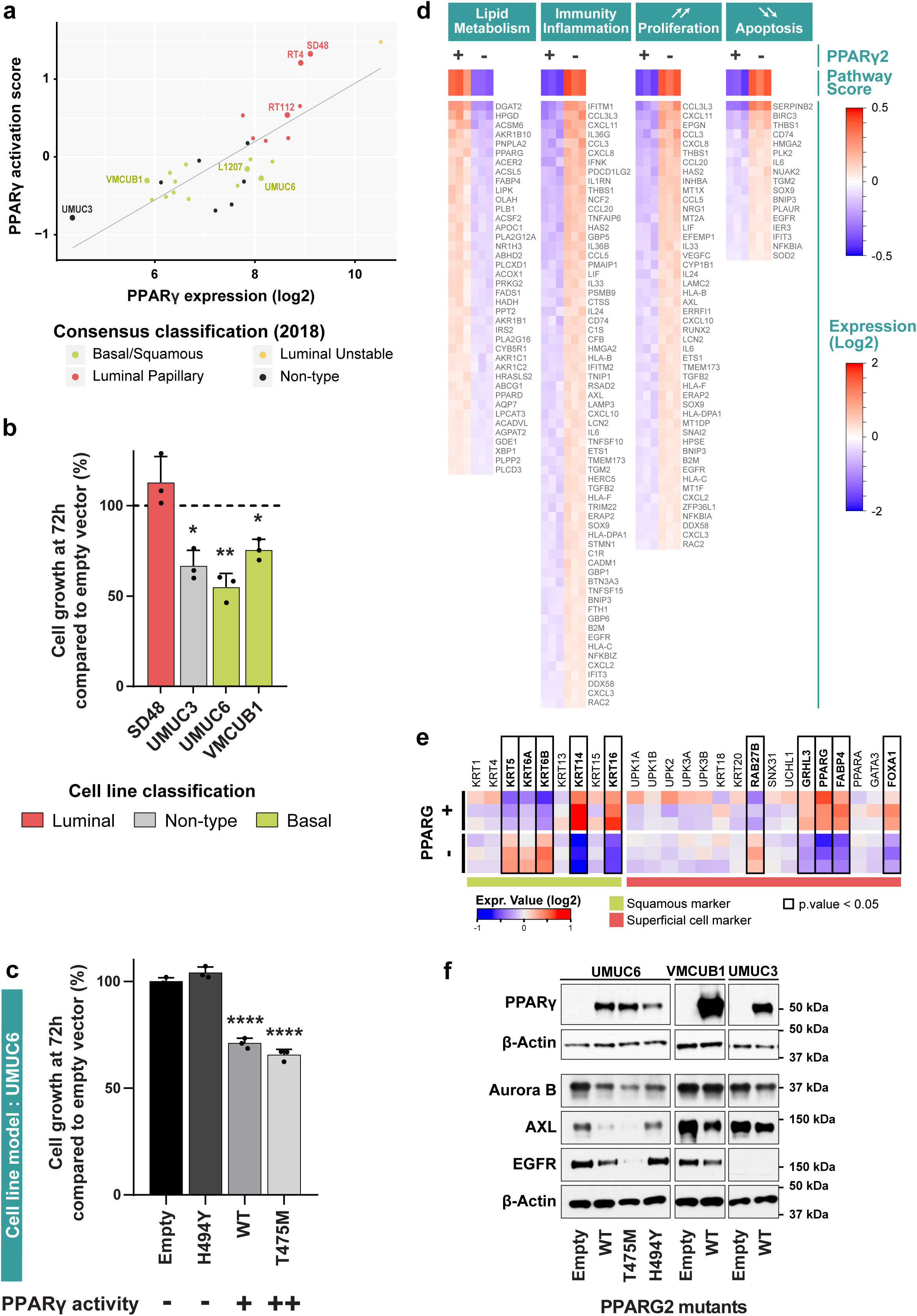
Effects of PPARγ overexpression on basal bladder cancer cell growth. **a)** PPARγ expression and activation score in a panel of bladder cancer cell lines. **b)** Four bladder cancer cell lines were transfected by a pRP vector encoding wild-type (WT) PPARγ. Expression of PPARγ was estimated after 48h by western-blot analysis (supplementary Fig.10c). The number of viable cells was quantified by CellTiter-Glo at 72h **c)** UMUC-6 cells were transfected with a pRP vector encoding wild-type (WT), inactive mutant (H494Y) or active mutant (T475M) PPARγ. Expression of PPARγ was estimated 48h later by western-blot analysis (supplementary Fig.10c). The number of viable cells was quantified by CellTiter-Glo at 72h. **b-c**: results with PPARγ (WT or mutant) were compared to those obtained with an empty vector using Dunnett’s multiple comparison test, * 0.01< *p*<0.05, ** 0.001< *p*<0.01, **** p<0,0001. **d-e)** Transcriptomic analysis upon PPARγ transient expression in UM-UC6 cells. PPARγ2 encoding plasmid (+), control backbone (-). DAVID analysis was performed for a list of 459 differentially expressed genes upon PPARγ overexpression (*p*-value < 0.05) to identify the biological process altered by PPARγ expression **(d)**. Expression of genes involved in urothelial differentiation is highlighted (**e). f)** Western-blot analysis of PPARγ, Aurora B, AXL and EGFR expression 72h after transient transfection of cells with a pRP vector encoding wild-type (WT), inactive mutant (H494Y) or active mutant (T475M) PPARγ. Actin was used as loading control.

### PPARγ LBD mutations F310S and H494Y favor an inactive conformation

To elucidate the structural basis for the deleterious functional effects of the mutations, we analyzed the crystal structures of PPARγ LBD F310S and H494Y (Supplementary Table 2). PPARγ F310 LBD mutant was crystallized in complex with GW1929 and PPARγ H494Y LBD in complex with GW1929 and the PGC1α coactivator peptide. Although the two mutants were less active than the wild-type protein, the ligand and/or coactivator peptide concentration used for crystallization allowed the proteins complexes to be crystallized in an active conformation. The GW1929 agonist ligand in the two mutant complexes maintained the same position as the WT complex forming similar interactions^16^ with the exception, in the F310S mutant, of the presence of two water molecules as a consequence of a larger binding pocket (Supplementary Fig. 9).

In the WT complex, H494 in helix 12 packs on top of V318 (H3) and forms intra-helical interactions with P495 and L496 that both contribute to a hydrophobic surface on which the coactivator packs (Fig. 4a). In the H494Y mutant, the side chain pointed toward loop 2-3 and the side interactions with P495 and L496 were lost suggesting a destabilization of helix 12. Due to the Y494 side chain re-orientation, Y494 forms new interactions with residues F315 (H3) and F292 and H294 (loop 2-3) of the mutant.

In the other mutant of interest, the bulky F310 side chain faces the loop 11-12 and is involved in van der Waals interactions with F388 (H7), I484 (H11) and M491 (loop 11-12). Because of the smaller size of the S310 side chain, these interactions are lost (Fig. 4b) and induce a more flexible conformation of the loop 11-12. In contrast S310 hydroxyl group interacts with the carbonyl moiety of Q311 and a water molecule. These variations in helix 11 and loop11-12 interactions will result in H12 destabilization and less efficient recruitment of coactivator. Overall, these data suggest that the two studied mutations, F310S and H494Y, are important residues for proper stabilization of helix 12 of PPARγ, preventing proper corepressor release and coactivator interaction.

### PPARγ displays an inhibitory effect on basal bladder cancer cells growth

The down-regulation of PPARγ in basal tumors suggested a potential tumor suppressive activity of PPARγ in these tumors. The association of PPARγ downregulation with PPARγ genomic deletions, DNA hypermethylation and of loss-of-function mutations, although rare events in basal tumors (observed in 4 out of 188 basal tumors, Fig. 2b), strongly reinforced this hypothesis. Activation of PPARγ by synthetic agonists has been a matter of debate regarding the PPARγ independent-effects of such molecules^20^. Therefore, to study the ability of PPARγ activation to inhibit basal bladder cancer cell proliferation, we transiently expressed PPARγ in three cell lines presenting a low PPARγ activation score and a low PPARγ expression level (UMUC-3, unclassified; UMUC-6 and VMCUB-1, Basal/Squamous cell lines), as well as in one luminal papillary cell line, SD48, which diplays a high expression level of PPARγ, a high PPARγ activation score and relies on PPARγ expression for its growth ^8^ (Fig. 5a and 5b). PPARγ overexpression, observed by western-blot or RT-qPCR analysis (Supplementary Fig. 10a and 10c), activated a PPARγ-dependent transcription program (Supplementary Fig. 10a) in the four cell lines. PPARγ overexpression induced a significant decrease in cell viability in the three cell lines expressing low levels of endogenous PPARγ, but did not affect the growth of SD48 cells (Figure 5b). Using UMUC-6 basal cells, we confirmed that the observed effects with the wild-type receptor or the activating mutation T475M were dependent on PPARγ activity since the overexpression of the inactive mutant of PPARγ, H494Y, had no effect neither on cell viability nor on PPARγ target gene expression (Fig. 5c and supplementary Fig. 10a-c). The tumor supressive role of PPARγ in basal tumors was further supported by our attempt to establish stable clones of basal UMUC-6 cells. We were only able to obtain a few clones with the wild-type or T475M PPARγ constructs, which turned out not to express PPARγ, whereas we obtained PPARγ-expressing clones using H494Y PPARγ construct (Supplementary Fig. 10d). To better understand the molecular mechanisms underlying this tumor suppressive property of PPARγ, we compared UMUC-6 transcriptomic data after transient transfection using either a control backbone or a PPARγ encoding plasmid. We identified 459 differentially expressed genes using LIMMA algorithm and considering a p-value<0.05 (Supplementary table 3). Analysis of the biological processes enriched in these genes using DAVID software highlighted that, as previously observed in luminal bladder cell lines, PPARγ expression induced an increased lipid metabolism^8,9,14^ and impaired immunity and inflammation^15^. However, PPARγ expression also induced a down-regulation of two sets of genes favoring cell proliferation and inhibiting apoptosis (Fig.5d). The regulation of these two process by PPARγ could account for the inhibition of cell viability induced upon PPARγ expression in basal cell lines. Focussing on the list of urothelial differentiation markers recently provided by Liu *et al.*^4^, we also confirmed the role of PPARG in inducing differentiation (Fig. 5e). However, as described by Warrick *et al.*^21^, PPARγ expression alone is not sufficient to transduce basal cells into luminal ones according to our consensus classifier for cell lines. We also performed a GSEA analysis of regulated genes using Reactome database which highlighted the up-regulation of “fatty acid metabolism” and ”fatty acid beta oxydation” as well as an increase of ”FOXO-mediated transcription of cell death genes”. A potential involvement of the EGFR pathway in the regulation of cell growth was suggested by the down-regulation of “GRB2 events in EGFR signaling”, which included a downregulation in gene expression of EGFR and its ligands EREG and EPGN (Supplementary Figure 11). We further validated this finding at the protein level by western-blot in UMUC-6, VMCUB1 and UMUC-3: PPARγ expression induced a down regulation of Aurora B, AXL and EGFR levels, which could regulate cell proliferation and apoptosis (Fig.5f). The down-regulations likely relied on PPARγ activity since they were stronger after the expression of the active mutant of PPARγ T475M but not observed after the expression of the inactive mutant PPARγ H494Y (Fig.5f, UMUC-6 cells).

## Discussion

Previous studies have suggested a tumor suppressor role for PPARγ in bladder cancer, based on its observed down-regulation in a subset of tumors and on the inhibitory effects of PPARγ agonist on bladder cancer cell lines *in vitro* and *in vivo* ^22–25^. Simvastatin-induced inhibition of bladder cancer cell growth was also attributed to the activation of the PPARγ pathway, further supporting its tumor suppressor role in bladder tumors^26^. In this study, we provided epigenetic, genetic and functionnal evidence to demonstrate the tumor suppressor role for PPARγ, and we associated this role to a particular subgroup of tumors, the basal subtype. In order to demonstrate its tumor suppressive properties, we overexpressed PPARγ in basal or unclassified cell lines expressing low levels of PPARγ, including UMUC-3 cells in which PPARγ expression induced comparable results to those observed recently using agonists^23^. These finding suggest that the observed effects using agonist molecules were more likely PPARγ-dependent. In addition, the other cell lines for which PPARγ agonists have been shown to induce cell growth inhibition ^22–25^ happened to be unclassified or basal using our consensus molecular classifier and thus supports the use of PPARγ synthetic agonists as a therapeutic option for basal tumors. The possible reactivation of PPARγ in basal tumors by agonists despite low expression levels in tumors presenting hemizygous deletions and/or promoter hypermethylation, and the fact that inactivating mutations did not seem to display dominant negative effects are in favor of an haplo-insufficiency of PPARγ. This mechanism of action of PPARγ has already been suggested in lipodystrophy^27^. A better understanding of the molecular basis of the tumor suppressive activity of PPARγ in basal bladder tumors may lay the groundwork to propose alternative therapeutic strategies to indirectly target the PPARγ pathway, in order to avoid various side effects induced by the available PPARγ synthetic agonists ^28,29^. We previsouly showed that basal tumors present an activation of the EGFR pathway and rely on EGFR activity for their growth *in vitro* and *in vivo*^18^. Here, we showed that PPARγ overexpression in basal cell lines induces a down-regulation of EGFR and its ligands, which was, at least for EGFR, dependent on PPARγ activity, since not observed upon the overexpression of the inactive mutant of PPARG, H494Y. These results suggest that the decreased cell viability induced by PPARγ overexpression may be partly EGFR-mediated. A tumor suppressor role for PPARγ has already been reported for several cancer types including colon, lung, breast and ovarian cancers, but the relationship with the EGFR pathway has not been reported in these cancers. So far, the tumor suppressive properties of PPARγ have been more linked to anti-angiogenic effects^30^ or to an increase in reactive oxygen species (ROS) level due to a metabolic switch induced by PPARγ^31^. The modulation of fatty acid metabolism and mitochondrial beta-oxydation by PPARγ in basal bladder cancer cell could also contribute to its tumor suppressive activities. *In vivo* studies should allow studying the impact of PPARγ expression on angiogenesis and the tumor microenvironment and their contribution on tumor growth. Inactivating mutations, altough not frequent, seem to be bladder cancer-specific, but deletions or methylation appear to be the main causes of PPARγ loss of activity. Inactivating mutations were initially reported in colon cancer but remain controversial since they have never been observed in independent cohorts^32^. A better knowledge of the structure/function effects of the loss-of-function mutation of PPARγ could also guide the design of new potent and more specific agonists.

Together with our previous studies that put forth the pro-tumorigenic role of PPARγ associated with DNA amplification and gain-of-function mutations^8,16^ in luminal tumors, this study highlights a dual role of PPARγ in bladder cancers. The therapeutic strategies targeting PPARγ should therefore be tumor subtype-dependent, strengthening the interest for the application of the molecular classification in the clinic. The dual role of PPARγ also suggest that the risk of bladder cancer associated with the use of pioglitazone^33,34^ should be associated with the development of luminal tumors. The cell context dependent effect of PPARγ has already been shown in breast and colon cancer^35–37^. A better understanding of the signaling pathways activated by the receptor to mediate both its pro-tumorigenic and tumor suppressive effects should help further our understanding of the relation between cell context, in particular cell differentiation, and PPARγ activity in bladder cancer but also in other tumor types.

## Methods

### Materials and chemicals

Rosiglitazone and GW1929 were purchased from Tocris Bioscience. The fluorescent PGC1α peptide (137-EAEEPSLLKKLLLAPA-152) and fluorescent NCoR1 peptide (2260-NLGLEDIIRKALMG-2273) were purchased from Thermo-Fisher. The PGC1α peptide (139-EEPSLLKKLLLAPA-152) and NCoR (2258-ASNLGLEDIIRKALMGS-2274) were synthesized by Pascal Eberling (IGBMC peptide synthesis common facility).

### Transcriptomic, genomic, methylation and ChIPseq data

We used transcriptomic data available for our CIT series of tumors^8,18^ and for 405 MIBC from TCGA (http://cancergenome.nih.gov). We used our affymetrix exon st.0 and U133 plus2.0 transcriptomic data^18^ available for RT112, L1207, VMCUB-1, UMUC-3, UMUC-6 cell lines. Tumors and cell lines were classified using a molecular consensus classification system^11^ and PPARG activation score were calculated as previously described taking into consideration the expression levels of 77 PPARG target genes^16^. We used publicly available copy number data for 402 MIBC from TCGA (http://cancergenome.nih.gov). We used methylation data (450k methylation array) available for 368 MIBC from TCGA and for 59 samples from our CIT series. We used histone marks (H3K27ac, H3K9me3) ChIPseq data for RT112, SD48 and L12017 and 5637 cell lines available in the GEO database: GSE104804 and GSE140891.

### Plasmid constructs

The pcDNA3-PPARγ2 and PPRE X3-TK-luc were generously provided by Pr. Chatterjee (Institute of Metabolic Science, IMS, Cambridge) and Bruce Spiegelman (Addgene plasmid #1015), respectively. We used pcDNA3.1-PPARγ2 and the QuikChange II Site-Directed Mutagenesis Kit (Agilent Technologies) according to the manufacturer’s protocol, to generate all the mutations. Mutations were confirmed by DNA sequencing. The GAL4 DNA-binding domain cloning vector pM and the activation-domain cloning vector pVP16 are part of the Mammalian Matchmaker Two-Hybrid Assay kit (BD Biosciences Clontech). The construct pM-MED1 (510-787) expressing the Gal4 DBD-MED1 nuclear receptor interacting domain was provided by Lieve Verlinden (KU Leuven, Belgium). The pCMX-GAL4N-SMRT was a gift of Makoto Makishima (Nihon University School of Medicine). The Gal4 DBD-NCoR1 NRID was kindly provided by William Bourget (Centre de Biochimie Structurale, Montpellier, France).

### Cell culture and transfection

The HEK293FT human cell line and the UMUC-6, UMUC-3, VMCUB-1, RT112 and 5637 human bladder tumor-derived cell lines were obtained from DSMZ (Heidelberg, Germany). HEK293FT, UMUC-6, UMUC-3, VMCUB-1, L1207 cells were cultured in DMEM, whereas 5637 and RT112 cells were cultured in RPMI. Media were supplemented with 10% fetal calf serum (FCS). Cells were incubated at 37°C, ^under an atmosphere containing 5% CO^2^. The identity of the cell lines used was checked^ by analyzing genomic alterations with comparative genomic hybridization arrays (CGH array), and FGFR3 and TP53 mutations were checked with the SNaPshot technique (for FGFR3) or by classical sequencing (for TP53). The results obtained were compared with the initial description of the cells. We routinely checked for mycoplasma contamination.

For reporter gene assays, HEK293FT cells were plated in 96-well plates (30,000 cells/well) and transfected with 30 ng pcDNA3-PPARγ2 (wild-type or mutated), 50 ng PPRE X3-TK-luc and 6 ng pRL-SV40 (Promega), in the presence of the Fugene HD transfection reagent (Promega), in accordance with the manufacturer’s protocol. Cells were stimulated with 1µM rosiglitazone 24 hours later. Luciferase activity was determined 24 hours later, with the Dual-Glo® Luciferase Assay System (Promega), according to the manufacturer’s instructions, and the results obtained were normalized with the *Renilla* luciferase signal obtained with the pRL-SV40 plasmid.

For PPARγ2 transient overexpression in the 5637 cell line, we used six-well plates, 250,000 cells seeded per well. These cells were transfected 24 h later with 2.5 µg of pcDNA3-PPARγ2 (wild-type or mutated) in the presence of the Fugene HD transfection reagent (Promega). PPARγ2 transient overexpression in the UMUC3, UMUC6, VMCUB1 and SD48 cell lines were performed in six-well plates, 500,000 cells were seeded per well for the UMUC3 cell line, and 250,000 cells for the three other cell lines. Cells were transfected 24 h later with 2.5 µg of pRP-PPARG2 wild-type or mutated vector (Vector Builder) in the presence of Fugene HD transection reagent according to manufacturer’s instructions (Promega). The cells were selected 24h after transfection with 4µg/mL of puromycin for 24h, and then seeded at respectively 7,000, 3,000, 3,000 and 2,500 cells per wells in ninety-six-well plates. Cell viability was assessed by CellTiter-Glo® Luminescent Cell Viability Assay (Promega) 72h later and normalized by the signal obtained just after plating.

RNA was extracted with the RNA easy mini kit (Qiagen) and proteins were extracted by cell lysis in Laemmli buffer (50 mM Tris-HCl (pH 7.5), 250 mM NaCl, 1% SDS) supplemented with protease inhibitors and phosphatase inhibitors (Roche) 48 h after transfection.

For mammalian two-hybrid assay, HEK293FT cells were plated in 96-well plates (30 000 cells/ well) and transfected with 20 ng pV16-PPARγ2 (wild-type or mutated), 20 ng pM-MED1, 50 ng pG5-luc (Promega) reporter plasmid and 6 ng pRL-SV40 (Promega), in the presence of the Fugene HD transfection reagent (Promega), in accordance with the manufacturer’s protocol. Luciferase activity was determined 48 hours later, with the Dual-Glo® Luciferase Assay System (Promega), according to the manufacturer’s instructions, and the results obtained were normalized with the Renilla luciferase signal obtained with the pRL-SV40 plasmid.

### Immunoblotting

Cell lysates were clarified by centrifugation. The protein concentration of the supernatants was determined with the BCA protein assay (Thermo Scientific). Ten µg of proteins were resolved by SDS-PAGE in 4-15% polyacrylamide gels, electrotransferred onto Biorad nitrocellulose membranes and incubated with primary antibodies against PPARγ (Abcam #ab41928, used at 1/1000) and β-actin (Sigma Aldrich #A2228, used at 1/25,000). Horseradish peroxidase-conjugated anti-mouse IgG (Cell Signaling Technology # 7074, used at 1/3,000) was used as the secondary antibody. Protein loading was checked by staining the membrane with Amido Black after electroblotting.

### Real-time reverse transcription-quantitative PCR

Reverse transcription was performed with 1 µg of total RNA, and a high-capacity cDNA reverse transcription kit (Applied Biosystems). cDNAs were amplified by PCR in a Roche real-time thermal cycler, with the Roche Taqman master mix (Roche) and Taqman probe/primer pairs that we previously used and described^16^. Relative gene expression was analyzed by the delta delta Ct method, with TBP as the reference.

### Biochemistry

The sequences encoding the ligand-binding domain of the His-hPPARγ (231-505) receptors was inserted into pET15b. Point mutations were introduced into *PPARγ* with the QuikChange II XL Site-Directed Mutagenesis kit (Agilent), in accordance with the manufacturer’s instructions.

The corresponding proteins were produced in *Escherichia coli* BL21 DE3 by overnight incubation at 22°C after induction with 1 mM IPTG at an OD_600_ of ~0.8. Soluble proteins were purified by Ni-NTA chromatography followed by size exclusion chromatography on a Superdex 200 (GE) column equilibrated in 20 mM Tris-HCl, pH 8.0, 200 mM NaCl, 5% glycerol, and 1 mM TCEP. The proteins were concentrated to 3-6 mg/mL with an Amicon Ultra 10 kDa MWCO. Purity and homogeneity of all proteins were assessed by SDS and Native Page (Supplementary Fig.2).

### Crystallization, X-ray data collection and crystal structure refinement

The crystallization experiments were performed by sitting drop vapor diffusion at 290 K, mixing equal volumes (200 nL) of protein at 5 mg/mL and reservoir solution. For all crystal structures, the data were indexed and integrated with XDS^38^ and scaled with AIMLESS^39,40^. The structure was solved by molecular replacement in PHASER^41^ and refined with PHENIX^42^ and BUSTER^43^ with TLS refinement, followed by iterative model building in COOT^44^.

Crystals of PPARγ F310S-GW1929 were grown in 25% PEG3350, 0.2 M LiSO4, BisTris 0.1M pH 6.5, transferred to artificial mother liquor containing 35% PEG3350 and flash-cooled in liquid nitrogen. X-ray diffraction data were collected at PX1 beamline of the SOLEIL synchrotron with a wavelength of 0.979 Å. The final structure was refined to Rwork and Rfree values of 16.8 and 20.6%, respectively, with excellent geometry (97.27% of residues in favored region of the Ramachandran plot and 2.73% in the allowed region). Crystals of PPARγ H494Y-GW1929-PGC1α were grown in 0.2 M ammonium acetate, 0.1 M Hepes pH 7.5, trisodium citrate 1.2M, transferred to artificial mother liquor containing 15% glycerol and flash-cooled in liquid nitrogen. X-ray diffraction data were collected at the ID30A3 beamline of ESRF with a wavelength of 0.968 Å. The final structure was refined to Rwork and Rfree values of 17.22 and 20.30%, respectively, with excellent geometry (97.81 % of residues in favored region of the Ramachandran plot and 2.19% in the allowed region). Data collection and refinement statistics are provided in Supplementary Table 1. GW1929 and side chains of the mutated residues of H494Y and F310S complexes could be modelled with confidence as shown into the Polder omit maps^45^ displaying reduced model bias and exclusion of solvent molecules (Supplementary Fig. 3b and 3c). All structural figures were prepared with PyMOL (www.pymol.org/).

### Microscale thermophoresis

measurements were performed with a Monolith NT.115 instrument (NanoTemper Technologies GmbH, Munchen, Germany). The PPARγ complexes were prepared in 20 mM Tris pH 8.0, 200 mM NaCl, 1 mM TCEP, 0.05% Tween 20. Each measurement consists of 16 reaction mixtures where the fluorescent-labeled peptide concentration was constant (70 nM) and serial dilutions of PPARγ LBD from a concentration of 100 μM down to 2 nM. Measurements were made with standard glass capillaries (Nanotemper) at 25°C, at 20-40% LED excitation and 80% MST power, with a laser-on time of 30 s and a laser-off time of 5s. NanoTemper Analysis 2.3 software was used to fit the data and to determine the K_d_.

### Thermal unfolding, nanoDSF

Fluorescence based thermal experiments were performed using Prometheus NT.48 (NanoTemper Technologies, Germany) with capillaries containing 10 μL PPARγ WT or mutants at 4mg/ml. The temperature was increased by a rate of 1 °C/min from 20 to 95 °C and the fluorescence at emission wavelengths of 330 nm and 350 nm was measured. NanoTemper PR.Stability Analysis v1.0.2 was used to fit the data and to determine the melting temperatures T_m_.

### Mass Spectrometry Analysis

Prior to mass spectrometry analysis, PPARγ and all the different mutant proteins were buffer exchanged against 200 mM of ammonium acetate at pH 6.8, using five cycles of concentration/dilution with a microconcentrator (Vivaspin, 10-KD cutoff, Sartorius, Göttingen, Germany). All the samples were diluted either in H2O/ACN/HCOOH (denaturing MS conditions) or in 200 mM AcONH4 (native MS conditions) to a final concentration of 5 µM and infused with an automated chip based nanoelectrospray device (Triversa Nanomate, Advion Bioscience, Ithaca, USA) operating in the positive ion mode, coupled to a Synapt G2 HDMS mass spectrometer (Waters, Manchester, UK).

### Affymetrix DNA array

In order to identify genes displaying changes in expression after PPARγ transient expression in UM-UC-6 cells, we transfected the cells for 96 hours with pRP-PPARG2 wild-type or backbone vector (Vector Builder). Three independent transfections were performed. mRNA was extracted and purified with RNeasy Mini kits (Qiagen). Total RNA (200 ng) from control and PPARγ expressing UM-UC6 cells was analyzed with the Affymetrix Human Clariom R DNA array. Raw gene expression data were normalized and summarized by the RMA (robust multi-array averaging) method (R package affy) with a customized chip definition developed by Microarray Lab, BrainArray (ClariomDHuman_Hs_ENTREZG_v22)^46,47^ The LIMMA algorithm was used to identify genes differentially expressed after PPARG expression. The *p*-values were adjusted for multiple testing by Benjamini–Hochberg FDR methods. Genes with a FDR below 5% were considered to be differentially expressed.

## Data availability

Atomic coordinates and related structure factors have been deposited in the Protein Data Bank with accession codes: 6T1S and 6T1V. Transcriptomic analysis of UMUC6 cell line following PPARγ2 overexpression have been deposited in the GEO database with accession codes GSE141230.

## Acknowledgements

This work was supported by a grant from Ligue Nationale Contre le Cancer (L.C.-T, H.N-K, C.K, J.F, C.DSG, F.D, E.C, Y.A., F.R., I.B.-P.) as an associated team (Equipe labellisée), the “Carte d’Identité des Tumeurs” program initiated, developed and funded by Ligue Nationale Contre le Cancer, by a “PL-Bio” project funded by INCa (2016–146), the French Ministry of Education and Research, the CNRS, and the Institut Curie. The project was also supported by the Ligue Régionale Grand-Est Contre le Cancer, the Centre National de la Recherche Scientifique, the Institut National de la Santé et de la Recherche Médicale and the University of Strasbourg. We acknowledge ANR-10-LABX-0030-INRT under the frame program Investissements d’Avenir labelled ANR-10-IDEX-0002-02ANR and the support and the use of resources of the French Infrastructure for Integrated Structural Biology FRISBI ANR-10-INBS-05, the Instruct-ERIC and the French Proteomic Infrastructure ProFI ANR-10-INBS-08–03. We would like to thank the staff of Proxima 1 at SOLEIL as well as of ID30A3 at ESRF for assistance in using the beamlines. We also thank Alastair McEwen (IGBMC) for help in X-ray data collections and Carole Peluso-Iltis for technical assistance. We would like to thank Simone Benhamou (IGR) and Thierry Lebret (Foch Hospital) for their help in setting up the CIT series of bladder tumors. We thank GIS IBiSA and Région Alsace for financial support in purchasing a Synapt G2 HDMS instrument.

## Author contributions

L.C-T, S.C, F.R, N.R. and I.B.-P. designed the study. F.R, N.R, S.C,Y.A, T.L and I.B.-P supervised the study. L.C.-T., H.N-K, J.F, C.DSG, E.C carried out the bioinformatics analysis. A.K and A. DR classified the tumors and cell lines according to the consensus molecular classification. L.C.-T., H.N-K, F.D., C.K. J.F. and I.B.-P. performed the functional studies and analyzed the data. S.B, Q.P and J.O. performed the biochemical, biophysical, and structural studies, and N.R. analyzed the data. M.L. performed the mass spectrometry experiments, M.L. and S.C. analyzed the mass spectrometry data. L.C.-T, H.N-K, J.F, N.R. and I.B.-P. wrote the manuscript. All authors made comments on the manuscript.

