## supplementary figures for "PPARγ is a tumor suppressor in basal bladder tumors offering new potential therapeutic opportunities"

**Supplementary Table 1**: **Clinical and pathological characteristics of TCGA and CIT tumors.**

| | TCGA | CIT Cohorte |
|---|---|---|
| **Total population** | 405 | 196 |
| **Sex** | | |
| Male | 299 | 156 |
| Female | 106 | 36 |
| NA | 0 | 4 |
| **Tumor stage** | | |
| Ta | 0 | 56 |
| T1 | 2 | 45 |
| T2 | 130 | 28 |
| T3 | 138 | 44 |
| T4 | 133 | 22 |
| NA | 2 | 1 |
| **Tumor classification** | | |
| LumP | 127 | 114 |
| LumU | 53 | 18 |
| LumNS | 20 | 11 |
| Stroma-rich | 45 | 13 |
| Ba/Sq | 152 | 36 |
| NE-like | 6 | 4 |
| NA | 2 | 0 |

### Supplementary Table 2: crystallographic data and refinement.

| | PPARγ2 F310S-GW1929 | PPARγ2 H494Y-GW1929-PGC1α |
|---|---|---|
| **PDB ID** | **6T1S** | **6T1V** |
| **Data collection** | | |
| **Space group** | $P4_12_12$ | I 2 2 2 |
| **Cell dimensions** | | |
| **a, b, c (Å)** | 60.66 60.66 169.52 | 56.35, 120.74, 149.84 |
| **a, b, g (°)** | 90, 90, 90 | 90, 90, 90 |
| **Resolution (Å)** | 49.33-1.65 | 52.74-2.21 |
| **Rmerge (%)** | 0.0096 (0.675) | 0.026 (0.291) |
| **Rpim (%)** | 0.0096 (0.675) | 0.026 (0.291) |
| **I/σI** | 25.77 (1.0) | 17.90 (2.43) |
| **Completeness (%)** | 99.72 (97.31) | 99.75 (99.96) |
| **Redundancy** | 2.0 (2.0) | 2.0. (2.0) |
| **CC(1/2)** | 1 (0.524) | 0.999 (0.842) |
| **Refinement** | | |
| **Resolution** | 49.33-1.65 | 56.35-2.21 |
| **N° reflections** | 39253 | 25889 |
| **Rwork/Rfree** | 0.168/0.206 | 0.184/0.198 |
| **N° atoms** | | |
| **Protein** | 2206 | 2250 |
| **Ligand/ion** | 52 | 37 |
| **Water** | 243 | 169 |
| **B-factors** | | |
| **Protein** | 48.34 | 48.46 |
| **Ligand/ion** | 50.88 | 36.02 |
| **Water** | 48.29 | 50.99 |
| **R.m.s. deviations** | | |
| **Bond lengths (Å)** | 0.016 | 0.004 |
| **Bond angles** | 1.74 | 0.98 |

**Supplementary Table 3**: **Up and down-regulated genes following PPARG2 overexpression**

| Gene | logFC | p_value | Gene | logFC | p_value | Gene | logFC | p_value | Gene | logFC | p_value |
|---|---|---|---|---|---|---|---|---|---|---|---|
| MMP19 | 2,0627 | 1,21E-03 | KRT16P1 | 0,8469 | 2,85E-02 | ENTPD3 | 0,6399 | 1,42E-02 | MAB21L3 | 0,5081 | 1,42E-02 |
| SLITRK6 | 1,9623 | 1,63E-03 | LOC647859 | 0,8462 | 1,86E-02 | CYFIP2 | 0,6372 | 4,51E-02 | ST6GALNAC4 | 0,5045 | 3,83E-02 |
| DGAT2 | 1,9461 | 1,21E-03 | PRR15 | 0,8457 | 4,98E-03 | TNFRSF21 | 0,6363 | 1,08E-02 | NR1D1 | 0,5044 | 2,23E-02 |
| SDC2 | 1,8178 | 5,62E-03 | SLC44A3 | 0,8306 | 5,64E-03 | PRKG2 | 0,6362 | 4,62E-02 | AKR1C2 | 0,5010 | 4,93E-02 |
| LOC100506253 | 1,7562 | 4,07E-02 | SERPINE1 | 0,8284 | 5,64E-03 | FRRS1 | 0,6337 | 3,36E-02 | HRASLS2 | 0,4936 | 3,28E-02 |
| C11orf86 | 1,6620 | 3,16E-03 | ACER2 | 0,8229 | 2,33E-02 | LOC107985544 | 0,6288 | 1,83E-02 | MIR1266 | 0,4916 | 2,71E-02 |
| AQP3 | 1,6444 | 3,19E-03 | B4GALT1 | 0,8184 | 5,86E-03 | DMBX1 | 0,6270 | 1,17E-02 | LOC101928635 | 0,4905 | 3,52E-02 |
| HPGD | 1,4955 | 1,29E-02 | TNFAIP2 | 0,8102 | 5,73E-03 | METTL7A | 0,6170 | 4,78E-02 | ABCG1 | 0,4884 | 2,23E-02 |
| HYAL4 | 1,4728 | 3,19E-03 | ACSL5 | 0,8075 | 9,89E-03 | FAM89A | 0,6161 | 1,85E-02 | PPARD | 0,4860 | 1,85E-02 |
| PRG4 | 1,4664 | 5,86E-03 | SNHG18 | 0,8036 | 3,88E-03 | TRIM31 | 0,6161 | 1,71E-02 | WSB2 | 0,4852 | 4,07E-02 |
| GPRIN3 | 1,4048 | 4,19E-03 | LYPD3 | 0,8015 | 3,88E-03 | FADS1 | 0,6100 | 3,36E-02 | S100A16 | 0,4851 | 2,87E-02 |
| LOC101927118 | 1,3952 | 3,88E-03 | FABP4 | 0,7930 | 6,78E-03 | SSPN | 0,6049 | 8,93E-03 | CORO2B | 0,4778 | 4,18E-02 |
| STAT4 | 1,3772 | 7,51E-03 | SPOCD1 | 0,7909 | 4,54E-03 | STEAP3 | 0,6011 | 1,86E-02 | ATP13A3 | 0,4770 | 3,65E-02 |
| PTGER4 | 1,3736 | 3,16E-03 | ABCG2 | 0,7856 | 2,79E-02 | DIO2 | 0,5997 | 1,29E-02 | ZNF750 | 0,4703 | 4,62E-02 |
| S100A9 | 1,3525 | 1,89E-02 | SMOX | 0,7842 | 7,67E-03 | FABP5P2 | 0,5979 | 3,86E-02 | MDFIC | 0,4702 | 3,98E-02 |
| KRT14 | 1,2979 | 2,54E-03 | LIPK | 0,7827 | 3,44E-02 | AKR1B1P7 | 0,5975 | 9,19E-03 | CSF1R | 0,4699 | 3,56E-02 |
| RHOF | 1,2973 | 4,42E-03 | S100A4 | 0,7671 | 3,70E-02 | ATP2B4 | 0,5914 | 3,46E-02 | UBE2Q2 | 0,4663 | 4,93E-02 |
| TGFBR3 | 1,2928 | 4,42E-03 | GMFG | 0,7643 | 7,56E-03 | HADH | 0,5912 | 2,85E-02 | SORBS1 | 0,4663 | 3,79E-02 |
| ACSM6 | 1,2830 | 4,98E-03 | IL1RAPL2 | 0,7611 | 2,78E-02 | BHLHE41 | 0,5869 | 1,75E-02 | LOC105377818 | 0,4655 | 2,74E-02 |
| SEMA5A | 1,2354 | 3,34E-03 | BBOX1 | 0,7579 | 2,00E-02 | PPT2 | 0,5831 | 1,15E-02 | CSRP2 | 0,4647 | 2,28E-02 |
| KLRC1 | 1,2334 | 2,12E-03 | LOC100421523 | 0,7518 | 3,88E-03 | HSPA6 | 0,5827 | 1,85E-02 | EPCAM | 0,4630 | 2,83E-02 |
| TGM5 | 1,2178 | 1,21E-03 | OLAH | 0,7452 | 2,42E-02 | SHC1P1 | 0,5805 | 1,65E-02 | C1orf220 | 0,4613 | 3,19E-02 |
| PADI4 | 1,2048 | 2,12E-03 | RNF128 | 0,7339 | 3,66E-02 | ABCC11 | 0,5771 | 1,14E-02 | AQP7 | 0,4602 | 4,18E-02 |
| BCL6 | 1,1894 | 4,62E-03 | LONRF3 | 0,7281 | 1,42E-02 | CSRP2P1 | 0,5746 | 1,85E-02 | PROM2 | 0,4542 | 2,72E-02 |
| SNORD123 | 1,1779 | 2,00E-02 | ETFDH | 0,7270 | 2,23E-02 | AKR1B1 | 0,5740 | 1,49E-02 | HSPA7 | 0,4538 | 3,52E-02 |
| AATBC | 1,1369 | 3,19E-03 | PLB1 | 0,7233 | 4,42E-03 | HEIH | 0,5718 | 2,07E-02 | SLC9A7P1 | 0,4516 | 4,31E-02 |
| LOC105369593 | 1,1238 | 2,37E-02 | EPHB6 | 0,7216 | 8,01E-03 | PHLDA3 | 0,5709 | 1,65E-02 | LPCAT3 | 0,4471 | 3,36E-02 |
| OR7E14P | 1,0806 | 1,51E-03 | LOC105374470 | 0,7105 | 2,72E-02 | IFRD1 | 0,5702 | 2,42E-02 | SLC45A3 | 0,4422 | 2,96E-02 |
| TIMP3 | 1,0538 | 2,12E-03 | ACKR3 | 0,7102 | 9,15E-03 | RDH10 | 0,5676 | 1,71E-02 | POU5F1B | 0,4377 | 2,45E-02 |
| AKR1B10 | 1,0483 | 1,03E-02 | SLC25A20 | 0,7073 | 2,53E-02 | LOC105369592 | 0,5642 | 2,08E-02 | ACADVL | 0,4363 | 2,87E-02 |
| RALGPS2 | 1,0430 | 1,40E-02 | ACSF2 | 0,7059 | 1,08E-02 | IRS2 | 0,5634 | 3,65E-02 | TMEM56 | 0,4352 | 4,75E-02 |
| PIM1 | 1,0035 | 3,88E-03 | TUFT1 | 0,7004 | 1,63E-02 | TFCP2L1 | 0,5612 | 3,33E-02 | GRHL3 | 0,4340 | 2,72E-02 |
| TRAJ23 | 0,9982 | 3,02E-02 | TTC39B | 0,6965 | 2,23E-02 | AKR1B1P3 | 0,5607 | 1,67E-02 | ITGAV | 0,4334 | 4,44E-02 |
| KRT16 | 0,9826 | 4,42E-03 | LBH | 0,6963 | 1,81E-02 | LMNTD1 | 0,5605 | 2,58E-02 | EPB41L1 | 0,4302 | 3,02E-02 |
| FAM213A | 0,9763 | 4,42E-03 | CXADR | 0,6906 | 2,79E-02 | ABHD6 | 0,5601 | 2,87E-02 | IMPDH1P4 | 0,4286 | 3,13E-02 |
| CEACAM1 | 0,9670 | 1,08E-02 | APOC1 | 0,6887 | 3,81E-02 | THRB | 0,5573 | 1,48E-02 | ARHGAP42P5 | 0,4280 | 4,66E-02 |
| AKR1B10P1 | 0,9616 | 4,98E-03 | LINC00847 | 0,6832 | 6,79E-03 | LOC102724908 | 0,5572 | 1,83E-02 | MARC1 | 0,4276 | 4,22E-02 |
| ZNF436 | 0,9541 | 3,16E-03 | EPAS1 | 0,6822 | 1,08E-02 | LOC105371688 | 0,5532 | 2,44E-02 | AGPAT2 | 0,4247 | 2,60E-02 |
| FOS | 0,9493 | 2,65E-02 | LRRC8B | 0,6764 | 4,65E-02 | HSPB8 | 0,5517 | 1,65E-02 | CDC26 | 0,4204 | 2,84E-02 |
| MFSD6 | 0,9454 | 2,48E-02 | ARHGAP26 | 0,6736 | 1,41E-02 | PLA2G16 | 0,5471 | 1,76E-02 | PTK2B | 0,4186 | 3,43E-02 |
| MEP1AP3 | 0,9439 | 2,14E-02 | CYP2C18 | 0,6682 | 3,98E-02 | C12orf45 | 0,5434 | 3,84E-02 | PLEKHA6 | 0,4185 | 3,02E-02 |
| DMBT1 | 0,9339 | 4,71E-02 | PLA2G12A | 0,6645 | 3,12E-02 | SLC4A11 | 0,5429 | 1,85E-02 | OAS1 | 0,4166 | 3,83E-02 |
| PNPLA2 | 0,9324 | 2,41E-03 | NR1H3 | 0,6643 | 8,78E-03 | AKR1C3 | 0,5406 | 4,39E-02 | FST | 0,4142 | 4,71E-02 |
| GOLGA7B | 0,9266 | 3,16E-03 | LOC102724608 | 0,6623 | 2,28E-02 | LOC105373998 | 0,5339 | 2,72E-02 | AKR1B15 | 0,4118 | 4,51E-02 |
| FOXO1 | 0,9178 | 1,10E-02 | ABHD2 | 0,6597 | 2,28E-02 | SHC1 | 0,5338 | 1,71E-02 | MAL2 | 0,4116 | 3,40E-02 |
| DENND2D | 0,9166 | 2,54E-03 | UAP1 | 0,6594 | 2,42E-02 | RHOD | 0,5335 | 1,77E-02 | POR | 0,4102 | 3,98E-02 |
| LOC107986632 | 0,8951 | 4,32E-02 | PLCXD1 | 0,6577 | 5,84E-03 | COL4A1 | 0,5329 | 1,63E-02 | H2AFJ | 0,4091 | 3,40E-02 |
| FAM213AP2 | 0,8913 | 1,29E-02 | HTRA1 | 0,6568 | 4,84E-02 | CYB5R1 | 0,5274 | 3,79E-02 | LOC724105 | 0,4079 | 4,55E-02 |
| S100A7 | 0,8903 | 2,36E-02 | ACOX1 | 0,6544 | 1,54E-02 | IL13RA2 | 0,5262 | 2,91E-02 | CDV3 | 0,4055 | 3,31E-02 |
| PPARG | 0,8894 | 1,18E-02 | S100A14 | 0,6506 | 3,40E-02 | DOCK8 | 0,5259 | 4,51E-02 | GDE1 | 0,4029 | 4,32E-02 |
| SPINK7 | 0,8751 | 3,56E-02 | LOC107986160 | 0,6496 | 3,46E-02 | LOC105373567 | 0,5253 | 4,81E-02 | PCSK6 | 0,4024 | 4,62E-02 |
| TLR4 | 0,8673 | 1,03E-02 | LINC01589 | 0,6450 | 1,09E-02 | ALAS1 | 0,5211 | 2,33E-02 | FOSL2 | 0,3986 | 4,22E-02 |
| LOC105376192 | 0,8638 | 3,02E-02 | LOC574538 | 0,6435 | 1,42E-02 | LINC01384 | 0,5189 | 3,70E-02 | XBP1 | 0,3892 | 4,32E-02 |
| FGFBP1 | 0,8614 | 4,18E-02 | CKMT1A | 0,6419 | 2,08E-02 | IMPDH1P6 | 0,5181 | 2,85E-02 | PLPP2 | 0,3880 | 5,00E-02 |
| ABLIM3 | 0,8573 | 4,54E-03 | EGLN1 | 0,6414 | 1,85E-02 | AKR1C1 | 0,5172 | 4,68E-02 | PPFIBP2 | 0,3850 | 4,62E-02 |
| LOC105372578 | 0,8551 | 1,81E-02 | PCED1B-AS1 | 0,6402 | 1,08E-02 | ARHGAP23 | 0,5106 | 2,28E-02 | PLCD3 | 0,3717 | 4,51E-02 |

| Gene | logFC | p_value | Gene | logFC | p_value | Gene | logFC | p_value | Gene | logFC | p_value |
|---|---|---|---|---|---|---|---|---|---|---|---|
| **SLC7A5** | -0,3772 | 4,22E-02 | **UNC5C** | -0,5136 | 2,00E-02 | **ETS1** | -0,6294 | 1,99E-02 | **CNN3** | -0,8165 | 2,87E-02 |
| **FRAS1** | -0,3778 | 4,09E-02 | **LINC00673** | -0,5142 | 3,39E-02 | **TNFSF10** | -0,6306 | 3,02E-02 | **NRP1** | -0,8177 | 4,62E-03 |
| **HPS3** | -0,3791 | 4,46E-02 | **MIR6748** | -0,5181 | 4,51E-02 | **DOCK10** | -0,6323 | 1,40E-02 | **RARRES3** | -0,8264 | 1,03E-02 |
| **RAC2** | -0,3799 | 4,68E-02 | **ELK3** | -0,5191 | 2,23E-02 | **IL6** | -0,6345 | 1,81E-02 | **RBMS2P1** | -0,8334 | 2,48E-02 |
| **GRB14** | -0,3800 | 4,78E-02 | **GBP6** | -0,5195 | 2,23E-02 | **PLK2** | -0,6367 | 2,46E-02 | **ANTXR2** | -0,8494 | 1,03E-02 |
| **LOC100506411** | -0,3800 | 4,09E-02 | **FTH1** | -0,5201 | 3,14E-02 | **LCN2** | -0,6376 | 1,29E-02 | **EFEMP1** | -0,8570 | 9,04E-03 |
| **TRIM21** | -0,3879 | 3,98E-02 | **PHLDB2** | -0,5205 | 2,28E-02 | **RUNX2** | -0,6402 | 7,04E-03 | **LIF** | -0,8625 | 6,10E-03 |
| **GPR63** | -0,3902 | 3,57E-02 | **BNIP3** | -0,5229 | 1,48E-02 | **ITGB8** | -0,6428 | 3,93E-02 | **MUC13** | -0,8767 | 4,91E-03 |
| **SOD2** | -0,3906 | 4,15E-02 | **TRAM2** | -0,5252 | 3,36E-02 | **CXCL10** | -0,6445 | 1,41E-02 | **STRIP2** | -0,8767 | 5,03E-03 |
| **LOC100131849** | -0,3923 | 4,46E-02 | **TNFSF15** | -0,5260 | 2,49E-02 | **ERRFI1** | -0,6477 | 1,65E-02 | **TOX** | -0,8782 | 1,08E-02 |
| **TPRA1** | -0,4037 | 3,61E-02 | **TENM2** | -0,5276 | 3,12E-02 | **TCN1** | -0,6478 | 4,22E-02 | **LOC107984462** | -0,8823 | 3,79E-02 |
| **LAMB2** | -0,4114 | 4,28E-02 | **ARSJ** | -0,5288 | 3,12E-02 | **LAMP3** | -0,6482 | 1,66E-02 | **PMAIP1** | -0,8907 | 7,56E-03 |
| **GJB5** | -0,4188 | 3,16E-02 | **HPSE** | -0,5307 | 2,91E-02 | **MT2P1** | -0,6496 | 1,40E-02 | **LOC541472** | -0,9098 | 3,51E-02 |
| **CXCL3** | -0,4264 | 4,02E-02 | **BTN3A3** | -0,5346 | 4,64E-02 | **SLC28A3** | -0,6497 | 6,79E-03 | **MT2A** | -0,9174 | 6,78E-03 |
| **ARNTL2** | -0,4330 | 4,02E-02 | **ADAMTS6** | -0,5348 | 4,62E-02 | **HERC3** | -0,6498 | 3,40E-02 | **NRG1** | -0,9327 | 2,13E-03 |
| **COL17A1** | -0,4371 | 4,64E-02 | **SYT14** | -0,5364 | 4,51E-02 | **IGFL2-AS1** | -0,6516 | 1,07E-02 | **SERPINB13** | -0,9353 | 1,85E-02 |
| **STARD5** | -0,4409 | 2,61E-02 | **GBP1** | -0,5389 | 1,71E-02 | **AXL** | -0,6537 | 1,40E-02 | **TMEM27** | -0,9536 | 6,78E-03 |
| **IRF6** | -0,4415 | 4,22E-02 | **LOC107986820** | -0,5471 | 1,98E-02 | **RSAD2** | -0,6537 | 3,20E-02 | **MYOSLID** | -0,9867 | 1,08E-02 |
| **ANXA8L1** | -0,4431 | 3,49E-02 | **RRAS** | -0,5497 | 1,29E-02 | **RTP4** | -0,6612 | 3,98E-02 | **EVI2B** | -1,0003 | 1,41E-02 |
| **DDX58** | -0,4434 | 2,28E-02 | **KYNU** | -0,5579 | 1,65E-02 | **PRR16** | -0,6688 | 1,14E-02 | **CCL5** | -1,0194 | 3,88E-03 |
| **CD68** | -0,4439 | 2,97E-02 | **NDRG1** | -0,5584 | 1,40E-02 | **TIPARP** | -0,6696 | 3,01E-02 | **MBOAT1** | -1,0281 | 1,03E-02 |
| **MYO19** | -0,4504 | 4,93E-02 | **CBLB** | -0,5596 | 1,49E-02 | **TNIP1** | -0,6702 | 5,78E-03 | **FDCSP** | -1,0334 | 2,23E-02 |
| **PLCG2** | -0,4520 | 3,26E-02 | **SERPINE2** | -0,5657 | 1,28E-02 | **DPYD** | -0,6715 | 4,91E-02 | **MT1X** | -1,0382 | 1,71E-03 |
| **WDR66** | -0,4529 | 2,51E-02 | **UBA7** | -0,5679 | 4,32E-02 | **IFITM2** | -0,6743 | 7,77E-03 | **IL36B** | -1,0530 | 1,95E-03 |
| **KLF7** | -0,4604 | 4,18E-02 | **KRT5** | -0,5682 | 4,22E-02 | **SQOR** | -0,6781 | 2,28E-02 | **LOC105374003** | -1,0648 | 7,51E-03 |
| **NFKBIA** | -0,4626 | 3,36E-02 | **SNAI2** | -0,5736 | 1,03E-02 | **LOC105372130** | -0,6827 | 3,02E-02 | **INHBA** | -1,0883 | 1,63E-03 |
| **ZFP36L1** | -0,4634 | 2,78E-02 | **NT5E** | -0,5767 | 1,17E-02 | **TGFBI** | -0,6968 | 2,08E-02 | **GBP5** | -1,0890 | 3,75E-03 |
| **SYNE2** | -0,4650 | 2,72E-02 | **LOC442309** | -0,5796 | 4,09E-02 | **ARL4C** | -0,6969 | 2,53E-02 | **HAS2** | -1,0900 | 5,86E-03 |
| **FHDC1** | -0,4662 | 4,22E-02 | **ADAMTS1** | -0,5802 | 1,63E-02 | **HLA-B** | -0,6988 | 4,54E-03 | **CDA** | -1,1039 | 7,51E-03 |
| **IFIT3** | -0,4679 | 3,36E-02 | **MT1DP** | -0,5833 | 1,42E-02 | **ADAM19** | -0,7075 | 7,67E-03 | **TNFAIP6** | -1,1039 | 2,54E-03 |
| **CXCL2** | -0,4681 | 3,10E-02 | **TMCC3** | -0,5834 | 1,29E-02 | **SLFN12** | -0,7109 | 3,47E-02 | **CCL20** | -1,1074 | 2,29E-03 |
| **GBP1P1** | -0,4703 | 2,48E-02 | **CADM1** | -0,5834 | 3,46E-02 | **NEFL** | -0,7110 | 1,75E-02 | **MMP13** | -1,1234 | 3,04E-02 |
| **ALDOC** | -0,4721 | 4,20E-02 | **EIF5A2P1** | -0,5852 | 1,63E-02 | **MSMB** | -0,7131 | 1,03E-02 | **LINC00704** | -1,1263 | 9,82E-03 |
| **EPB41L2** | -0,4754 | 2,85E-02 | **SLC2A3** | -0,5875 | 4,44E-02 | **F2RL2** | -0,7142 | 2,34E-02 | **UCA1** | -1,1278 | 2,54E-03 |
| **CPNE8** | -0,4776 | 4,68E-02 | **CAPRIN2** | -0,5882 | 1,71E-02 | **HMGA2** | -0,7153 | 4,98E-03 | **SERPINB7** | -1,1495 | 4,06E-03 |
| **HEG1** | -0,4778 | 4,62E-02 | **C1R** | -0,5893 | 1,17E-02 | **CFB** | -0,7158 | 1,03E-02 | **NCF2** | -1,1621 | 1,51E-03 |
| **MCOLN3** | -0,4813 | 1,85E-02 | **ANXA3** | -0,5900 | 1,85E-02 | **PDP1** | -0,7171 | 1,42E-02 | **THBS1** | -1,1629 | 2,12E-03 |
| **ANTXR1** | -0,4824 | 4,18E-02 | **STMN1** | -0,5919 | 2,23E-02 | **LOC107986951** | -0,7180 | 1,03E-02 | **IL1RN** | -1,2027 | 2,13E-03 |
| **B4GALT5** | -0,4841 | 2,84E-02 | **FAM160A1** | -0,5923 | 3,02E-02 | **LAMC2** | -0,7209 | 8,38E-03 | **SPTLC3** | -1,2268 | 1,09E-02 |
| **IER3** | -0,4882 | 2,64E-02 | **E2F7** | -0,5925 | 1,62E-02 | **C1S** | -0,7381 | 4,98E-03 | **SPRR2A** | -1,2456 | 3,18E-02 |
| **EXT1** | -0,4896 | 4,28E-02 | **HLA-DPA1** | -0,5968 | 2,10E-02 | **ATP1B1** | -0,7408 | 3,12E-02 | **GBP4** | -1,2649 | 3,60E-03 |
| **DCBLD1** | -0,4903 | 3,59E-02 | **SOX9** | -0,6029 | 1,65E-02 | **GALNT5** | -0,7504 | 6,30E-03 | **BIRC3** | -1,2690 | 2,12E-03 |
| **MT1F** | -0,4913 | 2,21E-02 | **TGFB1I1** | -0,6057 | 8,38E-03 | **DKK1** | -0,7538 | 3,57E-02 | **PDCD1LG2** | -1,2727 | 3,34E-03 |
| **P3H2** | -0,4968 | 2,23E-02 | **ERAP2** | -0,6075 | 4,28E-02 | **KRT6B** | -0,7538 | 2,28E-02 | **ALDH1A3** | -1,3208 | 2,12E-03 |
| **NFKBIZ** | -0,4971 | 2,44E-02 | **TRIM22** | -0,6084 | 2,87E-02 | **CD74** | -0,7600 | 2,68E-02 | **NAV3** | -1,3270 | 1,69E-03 |
| **HLA-C** | -0,4985 | 1,81E-02 | **TMEM2** | -0,6112 | 4,18E-02 | **IL24** | -0,7610 | 1,95E-02 | **IFNK** | -1,3308 | 4,91E-03 |
| **JARID2** | -0,4991 | 4,18E-02 | **HLA-F** | -0,6123 | 7,04E-03 | **GFPT2** | -0,7628 | 7,95E-03 | **HCP5** | -1,3367 | 3,75E-03 |
| **SERPINA1** | -0,4997 | 2,48E-02 | **CALB2** | -0,6153 | 1,68E-02 | **CTSS** | -0,7637 | 2,23E-02 | **CXCL8** | -1,3813 | 1,63E-03 |
| **LRRC49** | -0,5008 | 2,33E-02 | **TGFB2** | -0,6154 | 2,14E-02 | **PSMB9** | -0,7736 | 3,52E-03 | **CCL3** | -1,3858 | 1,08E-02 |
| **KRT81** | -0,5023 | 1,81E-02 | **HERC5** | -0,6159 | 1,85E-02 | **UNC13C** | -0,7747 | 1,85E-02 | **AMTN** | -1,4289 | 1,09E-02 |
| **BHLHE40** | -0,5027 | 2,83E-02 | **ZC2HC1A** | -0,6166 | 3,36E-02 | **LOC102723739** | -0,7766 | 1,69E-02 | **EPGN** | -1,4608 | 3,31E-03 |
| **EGFR** | -0,5031 | 2,34E-02 | **TGM2** | -0,6194 | 1,09E-02 | **CYP4V2** | -0,7826 | 4,07E-02 | **IL36G** | -1,5200 | 3,05E-03 |
| **SCN2A** | -0,5032 | 3,01E-02 | **SLC2A12** | -0,6211 | 3,03E-02 | **CYP1B1** | -0,7864 | 1,29E-02 | **CXCL11** | -1,6230 | 1,21E-03 |
| **COL12A1** | -0,5068 | 1,85E-02 | **NUAK2** | -0,6228 | 3,01E-02 | **PORCN** | -0,7893 | 1,03E-02 | **DUSP10** | -1,6256 | 1,21E-03 |
| **B2M** | -0,5074 | 2,00E-02 | **FENDRR** | -0,6273 | 3,12E-02 | **VEGFC** | -0,7974 | 6,79E-03 | **PTPRZ1** | -1,7074 | 2,12E-03 |
| **BNIP3P1** | -0,5075 | 2,23E-02 | **TMEM173** | -0,6274 | 1,03E-02 | **LOC105373426** | -0,8069 | 3,65E-02 | **CCL3L3** | -1,7724 | 4,62E-03 |
| **FBXL2** | -0,5079 | 2,33E-02 | **NME7** | -0,6276 | 2,87E-02 | **IL33** | -0,8093 | 2,12E-02 | **IFITM1** | -1,9225 | 1,51E-03 |
| **TM4SF1** | -0,5103 | 2,99E-02 | **LOC105376374** | -0,6279 | 3,12E-02 | **VSNL1** | -0,8146 | 4,19E-03 | **SERPINB2** | -2,4708 | 3,96E-04 |
| **PLAUR** | -0,5115 | 2,08E-02 | **RBMS2** | -0,6284 | 2,27E-02 | **SH3RF2** | -0,8153 | 3,19E-03 | | | |

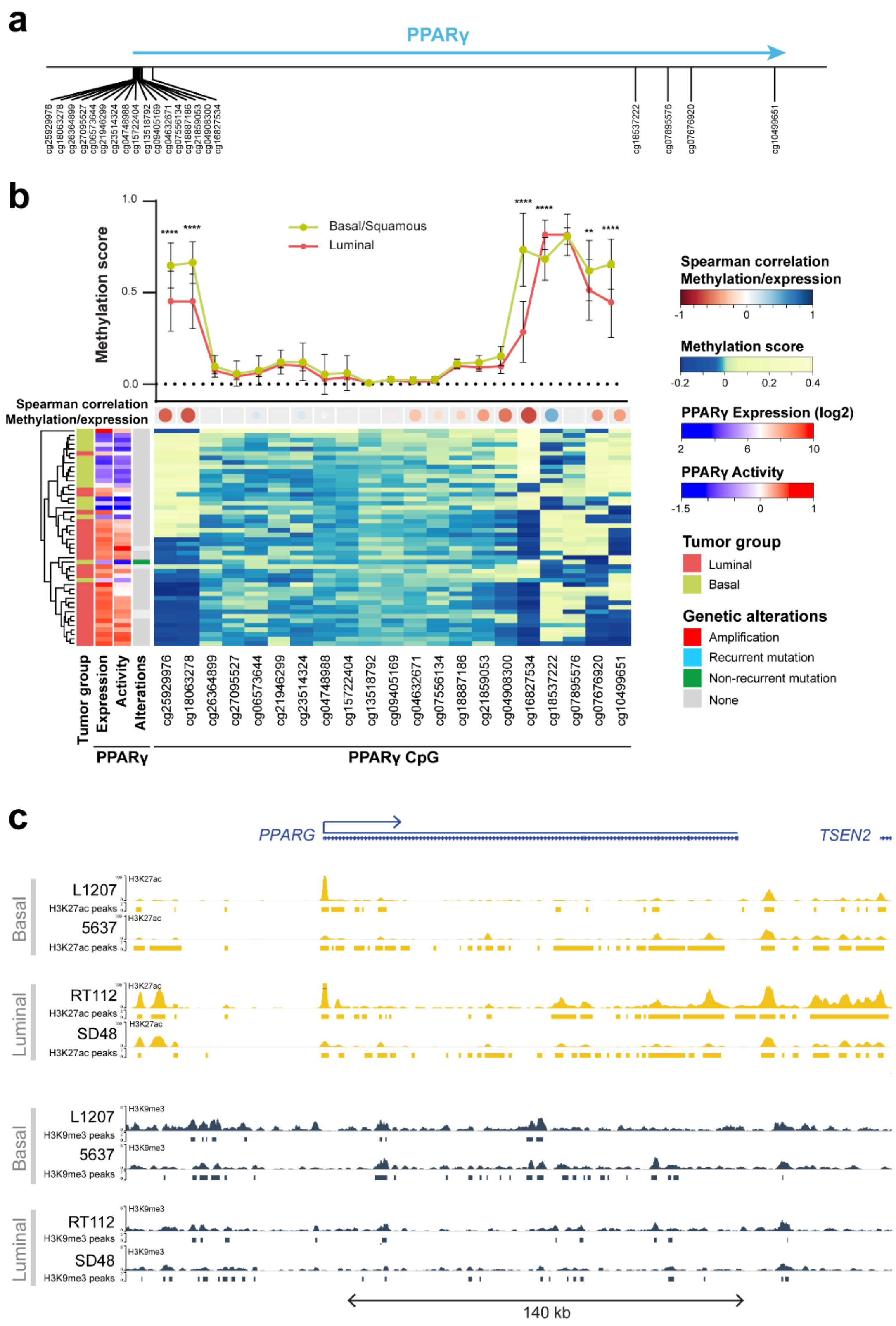


**Supplementary Figure 1: a)** Relative position of the CpG close to PPARG gene. **b)** Methylation of each CpG in CIT dataset for basal (green) and luminal (red) tumors (upper panel), correlation of CpG methylation and PPARG expression (middle panel) and heatmap representing centered methylation score for each tumor (lower panel).c**)** Genome Browser view of ChIPseq analysis for PPARg locus in Basal (L1207, 5637) and Luminal cell lines (RT112, SD48). ChIPseq aligned reads and associated peaks (MACS2) are represented in yellow and grey for H3K27ac and H3K9me3 marks respectively.

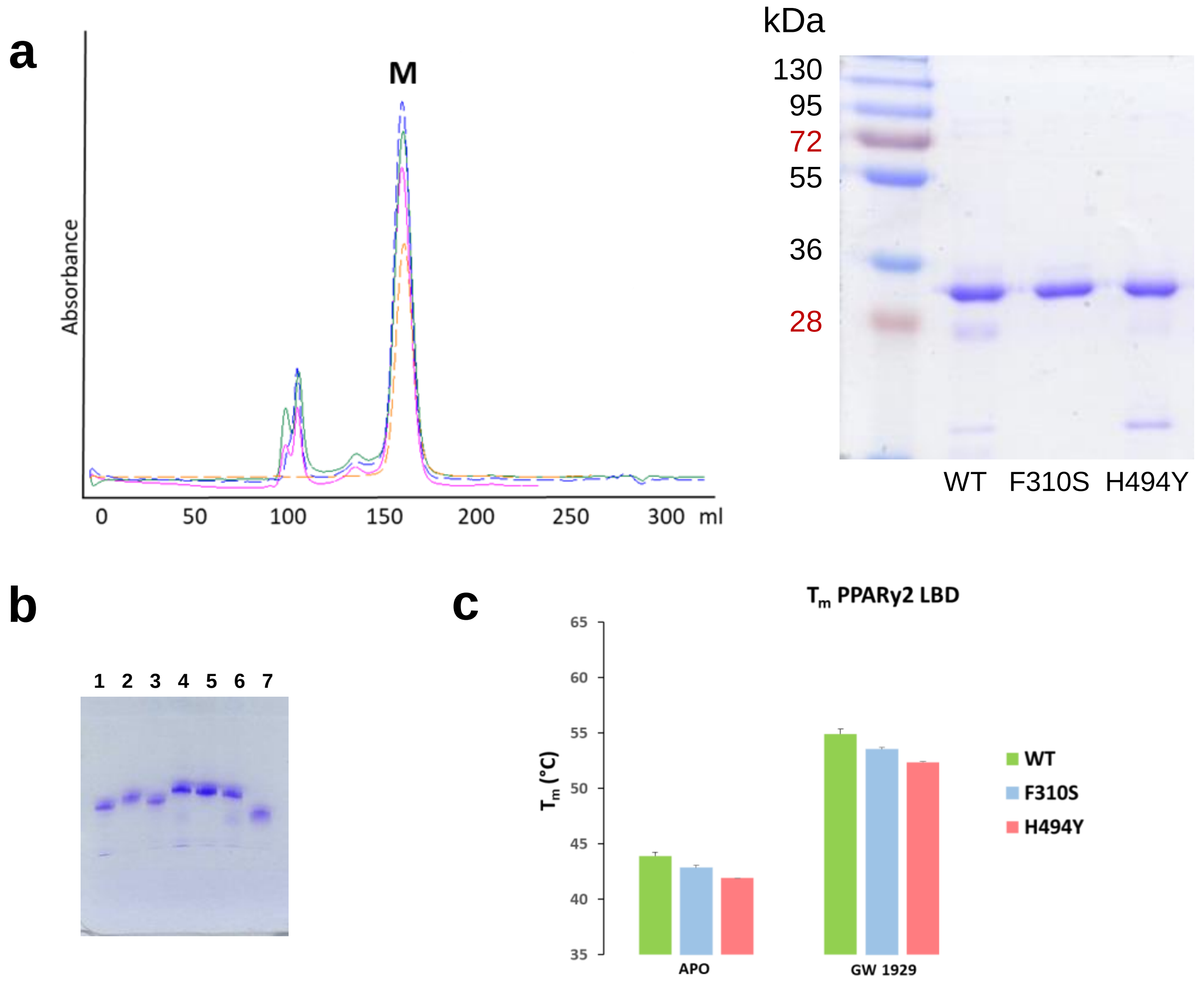


**Supplementary Figure 2:** Characterization of PPARγ mutants. **a)** Left: Gel filtration (S75 26/60) elution profile of apo PPARγ WT and mutants that eluate as monomers. Right: coomassie stained SDS gel corresponding to the PPARg wild-type (WT) and mutants monomeric peaks. **b)** Native PAGE of PPARγ WT (lane 1), F310S (lane 2), H494Y (lane 3), PPARγ WT with 1 equivalent of RXRα WT (lane 4), F310S with 1 equivalent of RXRα WT (lane 5), H494Y with1 equivalent of RXRα WT (lane 6) and RXRα monomeric WT (lane 7). **c)** Nano differential scanning fluorimetry comparing the thermal stability of the purified PPARγ WT and mutants alone and upon binding to the agonist ligand.

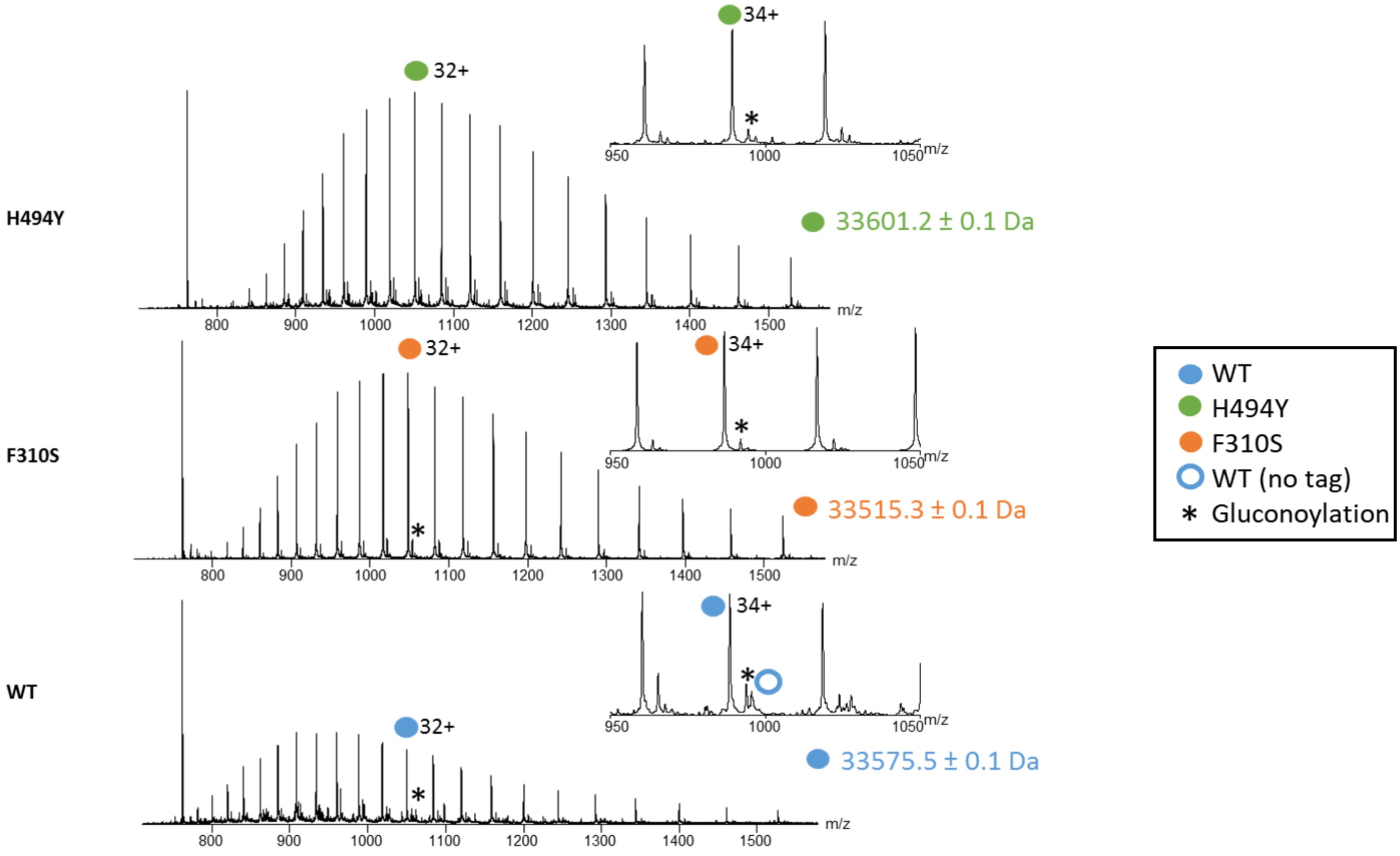


**Supplementary Figure 3:** Denaturing mass spectra of PPARγ LBD H494Y, F310S and WT. The experimental measured masses of the proteins are depicted in the right side of the mass spectra.

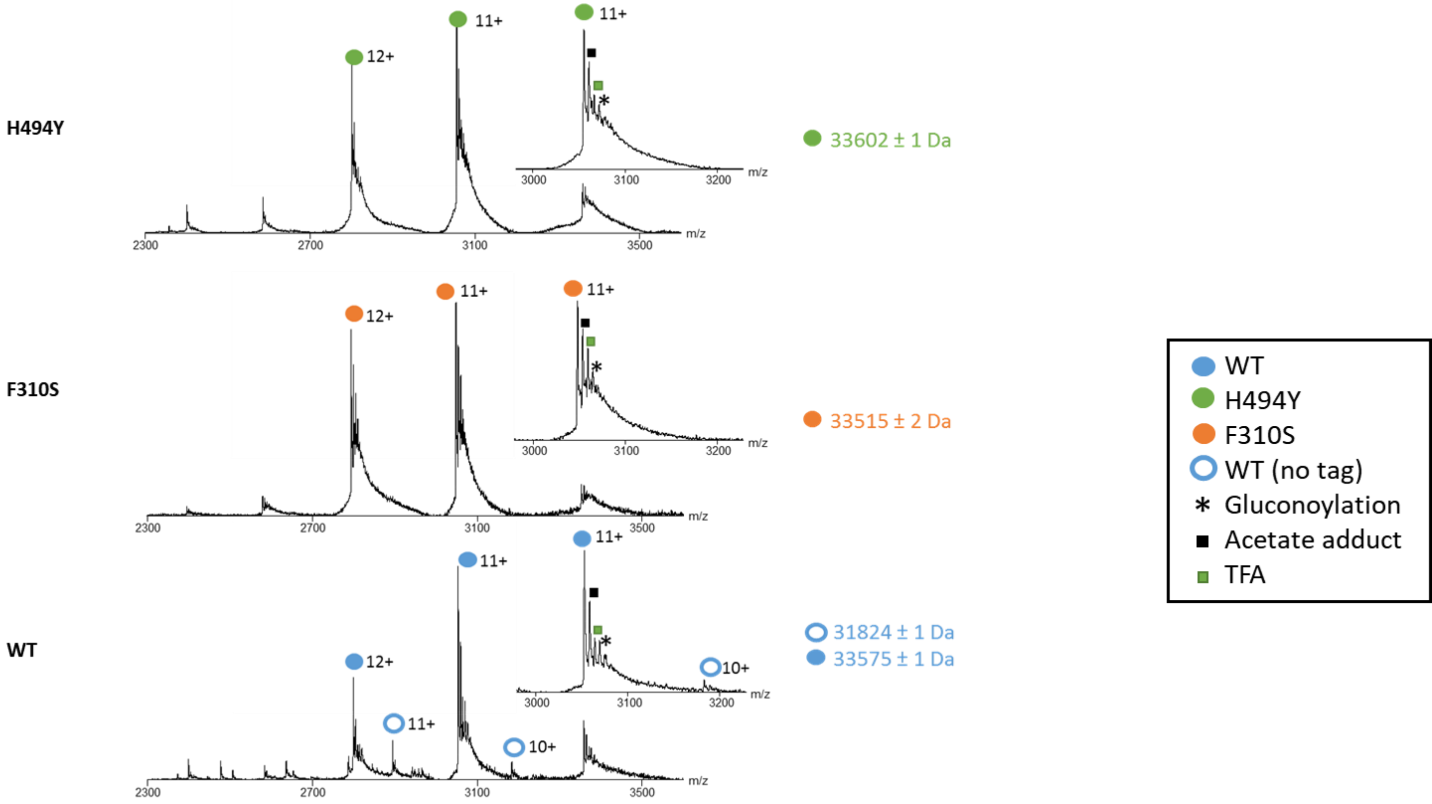


**Supplementary Figure 4:** Native electrospray ionization mass spectra of PPARγ LBD H494Y, F310S and WT. Mass spectra of apo proteins indicating the absence of any unexpected bound ligand in apo proteins.

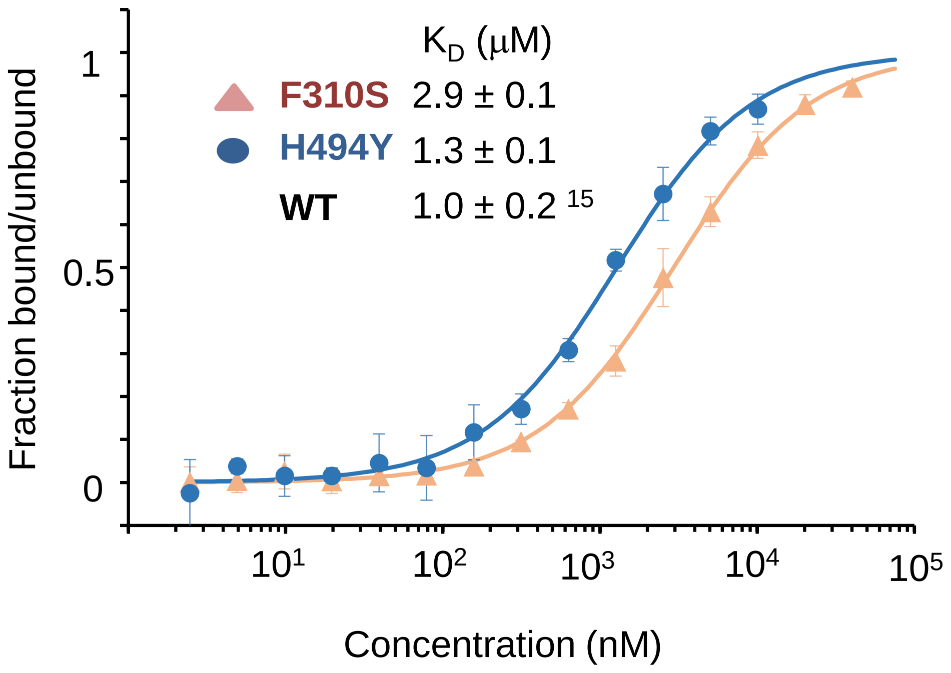


**Supplementary Figure 5:** Effect of PPARγ mutations on the PGC1α peptide interaction in presence of rosiglitazone as determined by microscale thermophoresis. Unlabeled PPARγ LBD protein was titrated into a fixed concentration of fluorescently labeled peptide in the presence of 3 equivalent of rosiglitazone. Isotherms were averaged over three consecutive measurements and fitted according to the law of mass action to yield the apparent $K_D$.

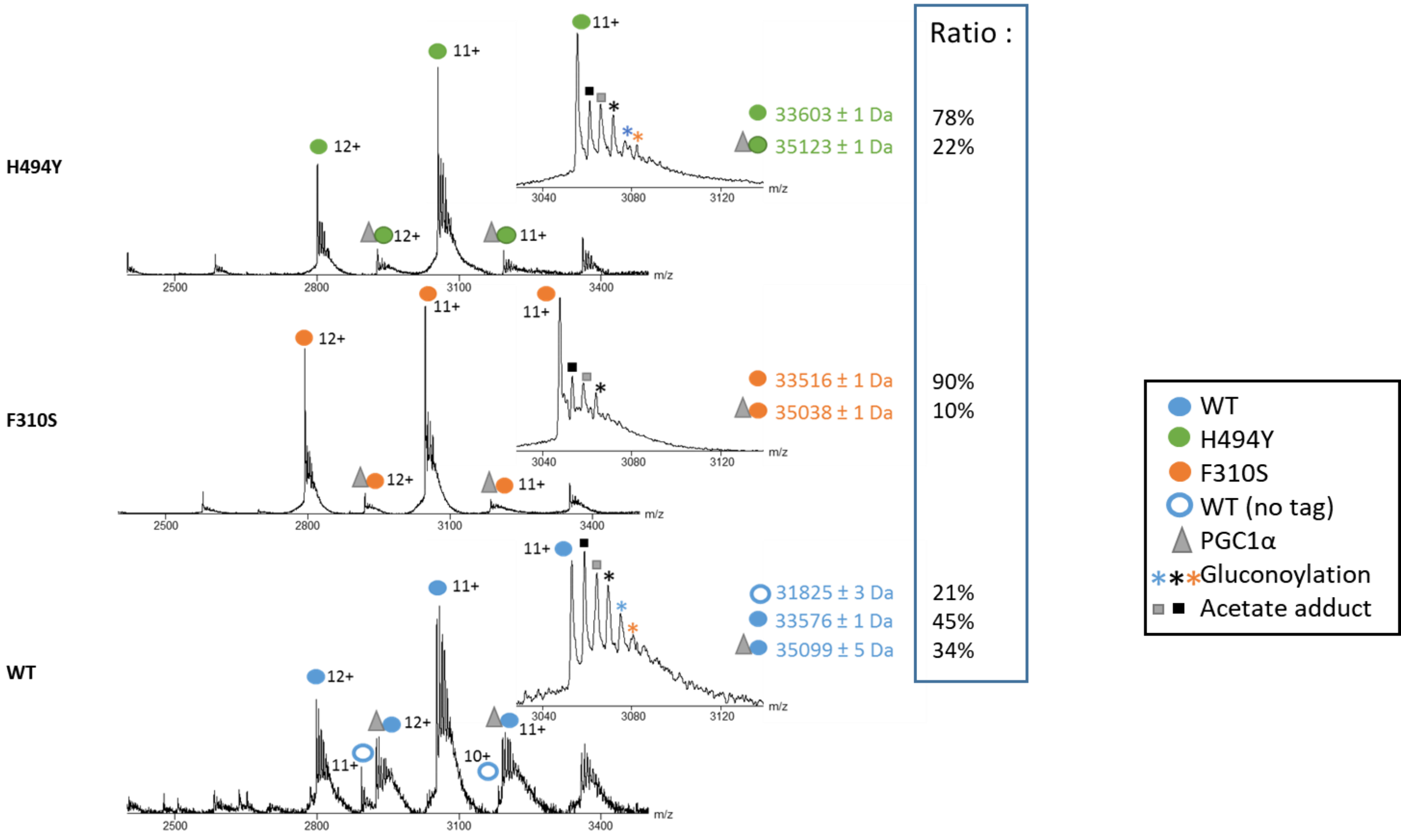


**Supplementary Figure 6:** Native electrospray ionization mass spectra of PPARγ LBD H494Y, F310S and WT. Mass spectra of the complexes obtained after addition of 3 fold excess of PGC1a coactivator peptide.

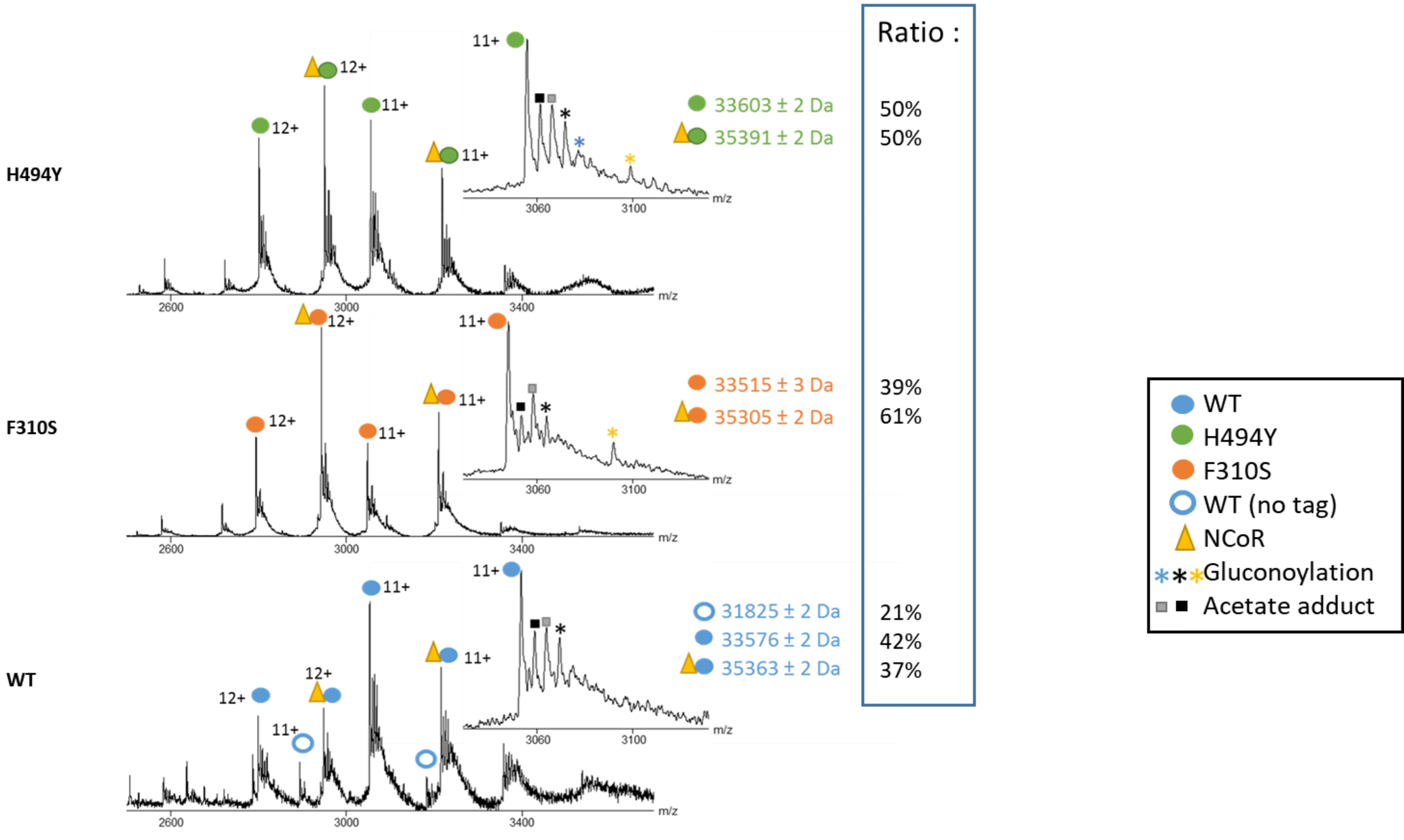


**Supplementary Figure 7:** Native electrospray ionization mass spectra of PPARγ LBD H494Y, F310S and WT. Mass spectra of the complexes obtained after addition of 5 fold excess of NCoR corepressor peptide

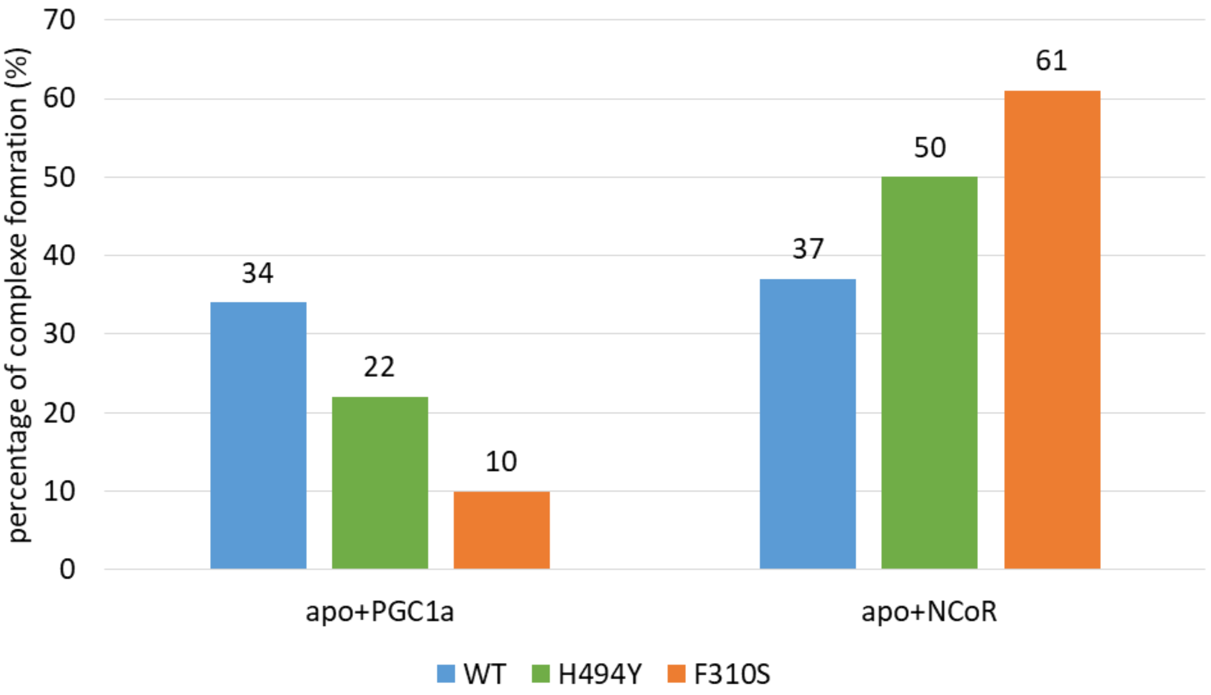


**Supplementary Figure 8:** Histogram of complexation rate between peptides PGC1α or NCoR and PPARg LBD H494Y, F310S and WT.

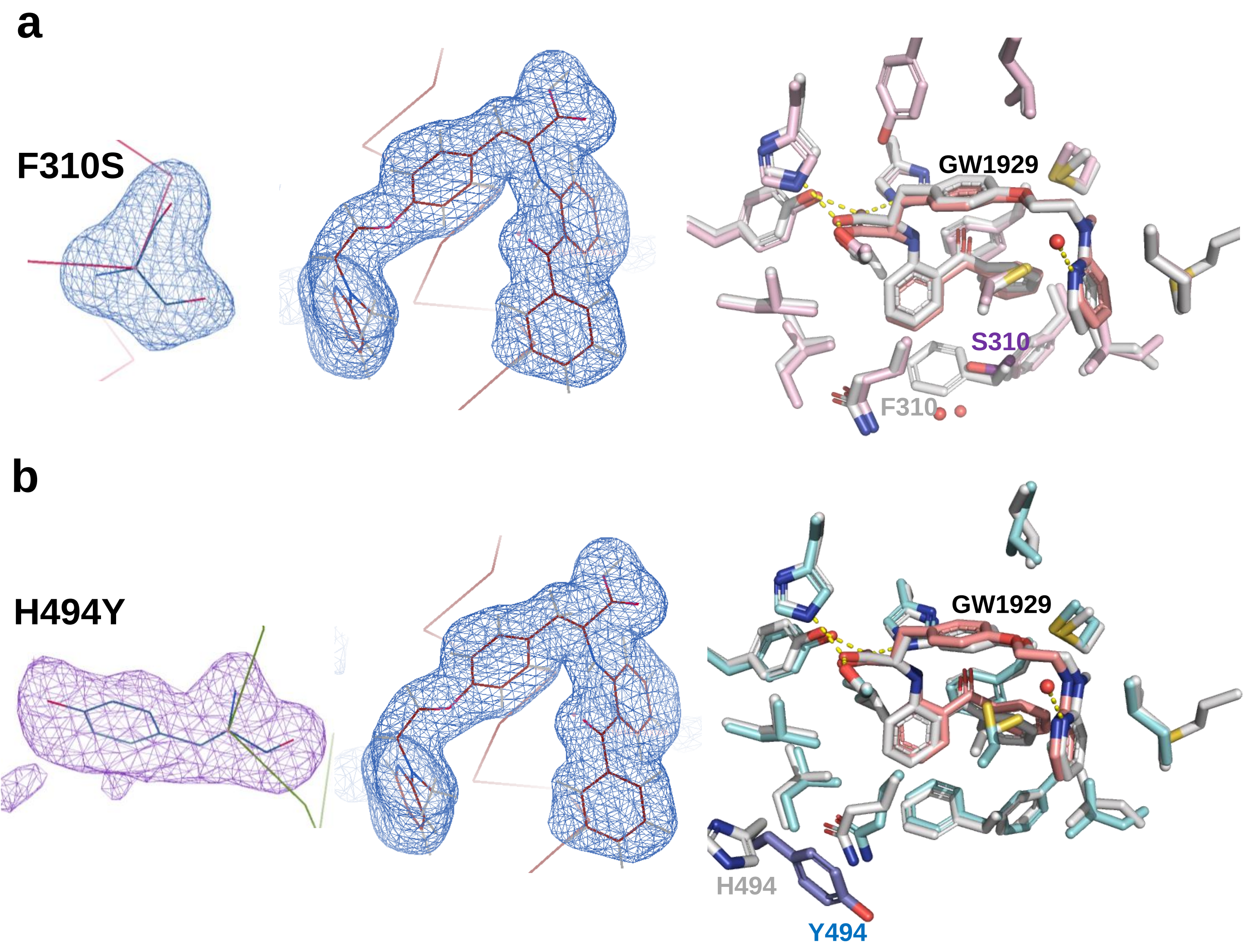


**Supplementary Figure 9:** GW1929 modelled into the difference density of the PPARγ **a)** F310S and **b)** H494Y crystal structures. Left: Residues mutated shown is an unbiased omit Polder map contoured at 3.5σ, with model bias reduction and exclusion of solvent density. Center: GW1929 modelled into the difference density of the mutants crystal structures. Shown is an unbiased omit Polder map contoured at 3.5σ, with model bias reduction and exclusion of solvent density. Right: Comparisons of the interactions made between GW1929 and residues lining the ligand binding pocket of PPARg mutants and WT (grey). Contributing side chains are shown as grey sticks, with residue forming hydrogen bonds (yellow dashed lines). Water molecules are shown by red spheres.

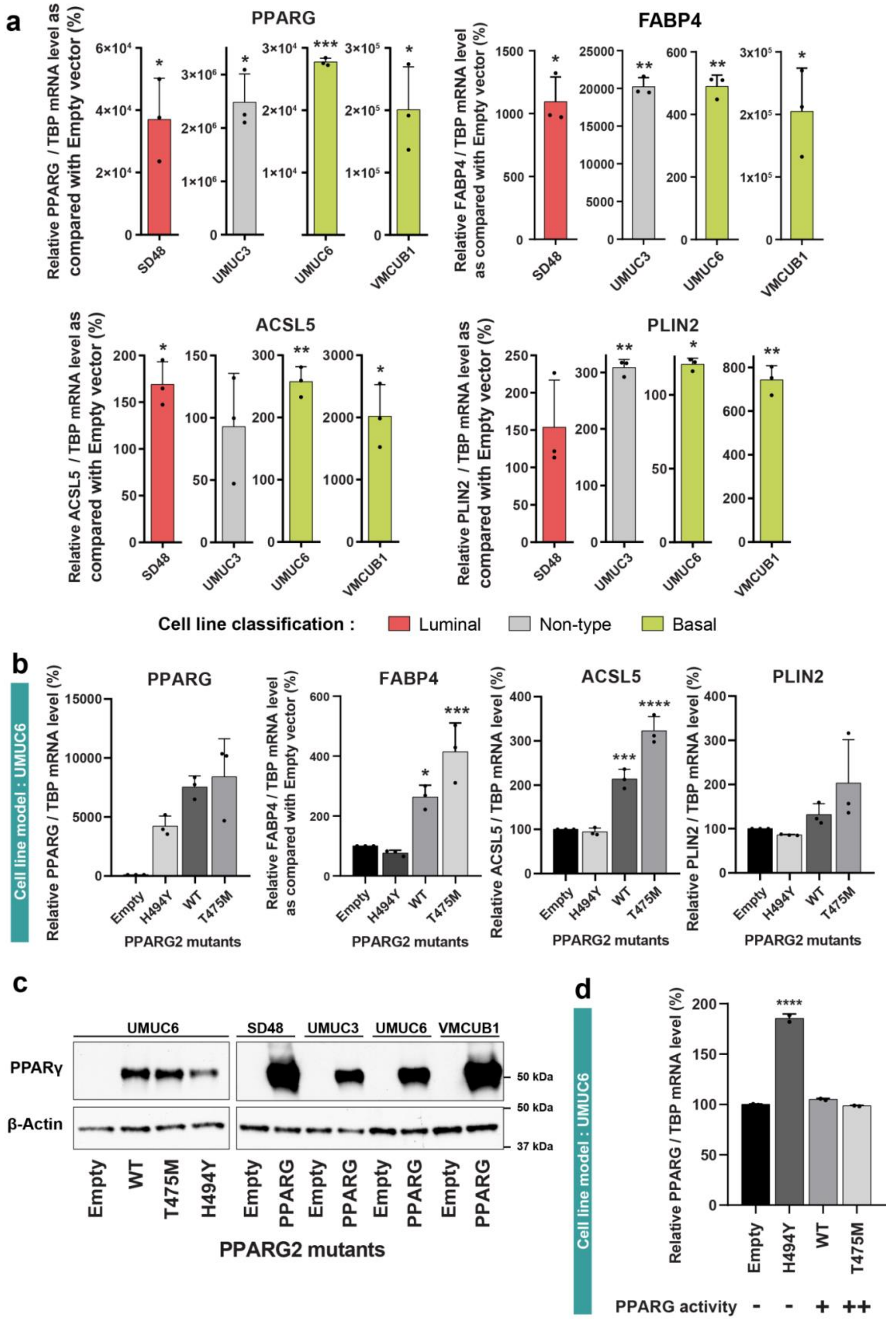


**Supplementary figure 10: Transient expression of PPARG induces a PPARg-dependent transcription in basal bladder cancer cell lines. a)** Expression of *PPARg* and *PPARg* -target genes 72 h after transfection by a pRP vector encoding *PPARg2* in 4 different bladder cancer cell lines. **b)** Expression of *PPARg* and *PPARg* target genes 72 h after transfection by a pRP vector encoding WT , gain-of-function (T475M) or loss-of-function mutant (H494Y) *PPARg2* in UMUC-6 basal bladder cancer cell lines. **c)** Expression of *PPARG* in pooled-UMUC6 cells transfected by a pRP vector encoding wt , gain-of-function (T475M) or loss-of-function mutant (H494Y) *PPARg2* . **a-c)** results obtained after PPARg transfection were compared to those obtained after transfection with the backbone vector using Dunnett's multiple comparison test, * 0.01< $p$<0.05, ** 0.001< $p$<0.01, **** $p$<0.0001 **d)**

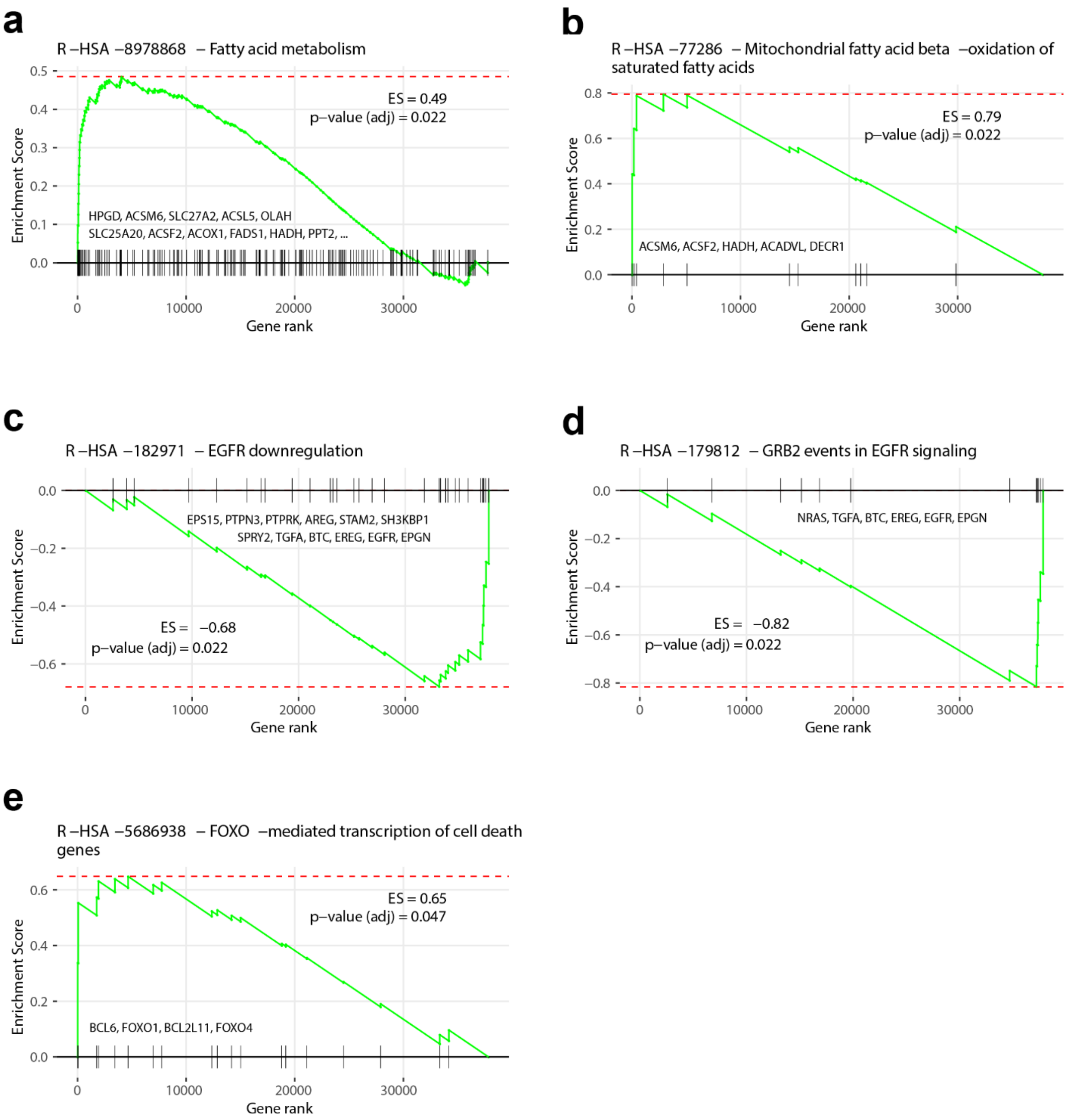


**Supplementary figure 11**: . Core enrichment genes, present in the leading or trailing edge, are annotated in the plot. Samples with overexpressed PPARγ were enriched with proteins involved in Fatty Acid Metabolism **(a)**, and, specifically, in Mitochondrial fatty acid beta−oxidation of saturated fatty acids **(b)**. Conversely, they presented EGFR downregulation **(c)** and were specifically depleted of GRB2 events in EGFR signaling **(d)**. Finally, after PPARγ overexpression, FOXO-mediated transcription of pro-apoptotic genes is upregulated **(e)**, which may contribute to the observed lower viability of cells.
